# Tit wit: environmental and genetic drivers of cognitive variation along an urbanization gradient

**DOI:** 10.1101/2024.10.30.621098

**Authors:** MJ Thompson, L Gervais, D Bharath, SP Caro, AS Chaine, C Perrier, D Réale, Anne Charmantier

**Affiliations:** Centre d’Ecologie Fonctionnelle et Evolutive, Univ Montpellier, CNRS, EPHE, IRD, Montpellier, France; Département des sciences biologiques, Université du Québec à Montréal, 141 Avenue du Président-Kennedy, Montréal, QC H2X 1Y4, Canada; Station d’Ecologie Théorique et Expérimentale du CNRS, UAR 2029, Moulis, France; UMR CBGP, INRAE, CIRAD, IRD, Institut Agro, Université Montpellier, Montpellier, France; Centro Agronómico Tropical de Investigación Y Enseñanza (CATIE), Turrialba, Costa Rica

**Author notes:** Joint first author (MJ Thompson, L Gervais).

**Keywords:** cognition, inhibitory control, common garden experiment, heritability, GWAS, great tit

## Abstract

Cognitive abilities can promote acclimation and adaptation to life in cities. However, the genetic versus environmental drivers of cognition have rarely been studied in the wild and there exists a major gap concerning the role of cognition in adaptation to novel urban contexts. To address this, we evaluate cognitive variation in wild great tits (*Parus major*; *N* = 393) along an urban gradient, and decipher the genetic basis of this variation using a combination of a common garden experiment, quantitative genetic analyses, and genome-wide association studies. Specifically, we measure inhibitory control abilities which affect how animals respond to novel resources and challenges. We find that wild urban and forest tits do not clearly differ in inhibitory control performance (number of errors or the latency to escape) during a motor detour task; a result that was consistent in birds from urban and forest origins reared in a common garden (*N* = 73).

Cognitive performance was repeatable (*R* = 0.35 – 0.38) and showed low to moderate heritability in the wild (*h^2^*= 0.16 - 0.28 using social and genomic pedigrees). We identified five SNPs that were significantly associated with the number of errors during the task, explaining 21% of the cognitive variation. These SNPs are linked to genes related to serotonergic and dopaminergic systems that are known to play important roles in cognition. Altogether, our study finds limited evidence that inhibitory control abilities have evolved under novel urban contexts, yet reveals a genetic basis of this cognitive trait in great tits.

## Introduction

Cities are expected to expand significantly in the coming decades (United Nations 2019).

This rapid expansion of urbanization is a process of novel environmental change that is transforming the habitat of wild populations worldwide by, for instance, increasing impervious surfaces, environmental pollution (chemical, light, sound), and introducing novel species and resources (Szulkin et al. 2020). Wild populations occupying urban environments need to adjust or adapt to these novel urban conditions to persist in these areas, and indeed phenotypic changes in urban populations are common and taxonomically widespread (Johnson and Munshi-South 2017; Diamond and Martin 2021; Lambert et al. 2021). For example, in comparison to their nonurban counterparts, some urban species are smaller (Hahs et al. 2023), less fearful of humans and associated stimuli (Geffroy et al. 2020), and behaviourally more aggressive (Miranda et al. 2013). Specifically, shifts in cognitive traits associated with the collection, storage, and use of environmental information could be especially important for facilitating adjustments to novelty and adaptation to urban ecological contexts (Sol et al. 2020; Lee and Thornton 2021).

Wild populations in cities are confronted with novel opportunities and challenges that are usually very different from those in natural environments, and successful urban organisms tend to be those that learn to exploit new resources and avoid harmful threats (Sol et al. 2013). There are several historical and modern examples of organisms innovating and exploiting new urban resources, including UK great tits (*Parus major*) opening glass milk bottles (Fisher and Hinde 1949) or Australian sulphur-crested cockatoos (*Cacatua galerita*) opening household waste bins (Klump et al. 2021). Some cognitive traits can reduce the risk of population extinction (Ducatez et al. 2020) and, specifically, learning, problem solving, and decision-making could be especially important for adjusting to novel conditions in cities (Griffin et al. 2017; Sol et al. 2020; Lee and Thornton 2021). Previous studies show that urban species and populations tend to have higher cognitive performance, especially on innovation, problem solving, or foraging related tasks, but high heterogeneity in results among the few existing studies prohibits generalizations (Sol et al. 2020; Vincze and Kovács 2022). At present, however, a fundamental gap exists on the potential for cognitive traits to evolve in wild populations.

Evaluating the evolutionary potential of cognitive traits requires examinations of the fitness consequences and genetic bases of these traits (Cauchoix and Chaine 2016; Morand-Ferron et al. 2016). In wild birds, higher cognitive performance on reversal learning, problem solving, and spatial memory tasks can be associated with both fitness benefits (Cauchard et al. 2012; Cole et al. 2012; Sonnenberg et al. 2019) and costs (i.e., problem solving and reversal learning; Cole et al. 2012; Madden et al. 2018). Evolutionary change in response to natural selection requires phenotypic variation to comprise underlying genetic variation and, to date, most estimates of heritability (i.e. proportion of phenotypic variance due to additive genetic variance) in cognition come from human or captive populations (Croston et al. 2015). Few studies have examined the heritability of cognitive traits in wild populations and, so far, heritability estimates range from high (Branch et al. 2022) to low (Quinn et al. 2016; De Meester et al. 2022; McCallum and Shaw 2023; Van Den Heuvel et al. 2023; Speechley et al. 2024; see also Sorato et al. 2018; Langley et al. 2020). For example, wild toutouwai (North Island robin; *Petroica longipes*) in New Zealand showed repeatable individual differences in inhibitory control (i.e., the ability to inhibit prepotent responses) over a year period, but the estimated heritability of this cognitive trait was low and did not differ from zero, suggesting this trait is strongly shaped by environmental effects (McCallum and Shaw 2023).

Evolutionary change is not solely dependent on heritability; it also hinges on the genetic architecture, which includes the number, magnitude, and physical associations of genes (Mackay 2001; Yamamichi 2022). Integrating information on the genetic architecture of cognitive traits with heritability estimates will, thus, allow more comprehensive predictions of evolutionary responses. For example, the speed of a trait’s response to selection can vary significantly depending on whether the trait is governed by a few genes with large effects (oligogenic = generally slower evolutionary response) or by many genes with small effects (polygenic = generally faster evolutionary response; Jain and Stephan 2017; Barghi et al. 2019). Few studies have examined the genetic architecture of cognition in wild populations. However, one study found significant associations between spatial memory and several genes involved in neuron growth and development in wild mountain chickadees (*Poecile gambelii*; Branch et al. 2022; Semenov et al. 2024). Recently, the authors found more than 95 genes correlated with spatial memory in this species, suggesting this cognitive trait might have a polygenic basis (Semenov et al. 2024). Despite growing evidence that the cognitive traits of urban organisms differ from their nonurban conspecifics (Sol et al. 2020; Lee and Thornton 2021), no studies have evaluated the evolutionary potential of cognition in urban populations. Recently, Sol et al. (2020) called for more studies that examine cognition under an urban evolution lens and, specifically, highlighted the usefulness of studies that disentangle the genetic and environmental drivers of cognition.

We first aimed to compare inhibitory control between wild urban and forest populations of great tits (*Parus major*) in and around the city of Montpellier, France using a motor detour task that is easily administered in the field (aim 1). Inhibitory control, or the ability to inhibit prepotent responses (MacLean et al. 2014; Kabadayi et al. 2018), is a cognitive trait that has been suggested to affect several fitness-related behaviours including foraging flexibility for novel resources (Coomes et al. 2021), dietary breadth (van Horik et al. 2018), premature attack on prey (Miller et al. 2019), conspecific resource sharing (MacLean et al. 2014), or innovation and problem solving (Lee and Thornton 2021). The ability to inhibit predominant responses when confronted with novel resources or challenges could play an important role in how organisms adjust to urban environments (Lee and Thornton 2021), but currently no studies have compared how these abilities differ between wild urban and nonurban populations. Since urban animals tend to have higher cognitive performance than their nonurban conspecifics (e.g., problem solving; Vincze and Kovács 2022), we hypothesize that wild urban great tits will have higher inhibitory control than forest tits. More specifically, we predict that wild urban great tits will make fewer errors and reach the goal more quickly in a motor detour task presented in the field.

Our second aim was to evaluate the genetic basis of inhibitory control and we used three complementary approaches to achieve this: a quantitative genetic analysis, a common garden experiment, and a Genome Wide Association Study (GWAS; aim 2). Recent studies on the genetic basis and evolutionary potential of inhibitory control in wild or recently descended populations have found low to moderate heritability estimates for this trait (e.g., Langley et al. 2020; McCallum and Shaw 2023; Prentice et al. 2023). Environmental variation can also affect inhibitory control where, for example, spatially unpredictable environmental conditions during development increase inhibitory control performance (van Horik et al. 2019) and heat stress and traffic noise decrease performance (Blackburn et al. 2022; Templeton et al. 2023; Soravia et al. 2023). To estimate the heritability of inhibitory control in the wild, we first used a quantitative genetic analysis combining genomic data and pedigrees from wild great tit populations. Since previous studies show inhibitory control may have some genetic basis, we predicted that inhibitory control would have underlying genetic variation and, thus, be heritable in the wild.

Second, to test for genetic differentiation in inhibitory control between wild urban and forest birds, we reared urban and forest great tits from eggs under the same environmental conditions. Common gardens have revealed genetic differences between urban and nonurban populations for physiological, morphological, and behavioural traits (e.g., Partecke et al. 2006; Winchell et al. 2016; Brans et al. 2018; Diamond et al. 2018), but urban cognition is yet to be examined in common garden experiments. In line with the expected differences in the wild, we predicted that tits from urban origins would have higher performance on the motor detour task (i.e., fewer errors and faster latency to goal) than tits from forest origins. Genetic and plastic changes can act in opposite directions to hide phenotypic divergences between wild populations (i.e., countergradient variation; Conover et al. 2009) and so, if urban and forest great tits do not clearly differ in their inhibitory control, we wanted to confirm the absence of genetic differences in this cognitive trait between populations using the common garden. Third, we used GWAS to identify candidate areas of the genome that associate with inhibitory control variation in the wild. Given that cognitive abilities are influenced by neurotransmitters such as dopamine or serotonin (Durstewitz et al. 1999) and neural growth (Kempermann et al. 2004; Barnea and Pravosudov 2011), we expected to identify genes associated with the nervous system. Further, we hypothesized that because inhibitory control is a quantitative trait it should have, like most quantitative traits, a polygenic basis (Mackay 2001; Flint 2003; Croston et al. 2015), likely involving many genes with small individual effects.

## Methods

### Study system

We monitored the reproduction of great tits at nest boxes in and around the city of Montpellier as a part of a long-term study (Charmantier et al. 2017). The forest population is located 20km north of Montpellier in La Rouvière forest (monitoring initiated in 1991; between 37 – 119 nest boxes with 32 mm diameter entrance, suited for both blue and great tits) and the urban population is monitored across eight study sites throughout the city of Montpellier that differ in their degree of urbanization (monitoring initiated in 2011; total 163 – 208 nest boxes). Fluctuations in nest boxes between years was a result of either theft or altering the proportion of large and small diameter nest boxes (32 vs 28 mm) to accommodate changing research objectives; the latter being more common in La Rouvière forest where blue tits are continuously monitored. During the Spring breeding season, we visited each nest box weekly to follow the reproduction of tits. When nestlings were approximately 12 days old, we caught parents in the nest boxes to ring them with a unique metal band, age them based on plumage (adult: 2+ year old, juvenile: 1 year old), take a blood sample, and measure several morphological and behavioural phenotypes (for more information see Caizergues et al. 2018; Caizergues et al. 2022). During the 2021 – 2023 breeding seasons, we also assayed individuals on a cognitive task in the field (see section ‘Cognitive assay’ below).

We quantified the level of urbanization at each study site using the proportion of impervious surface area (ISA; sealed non-natural surfaces including sidewalks, roads, and buildings). We used the imperviousness density data set from the Copernicus online database (resolution 10m, tiles: E38N22/E38N23, projection: LAEA EPSG 3035; European Environment Agency 2020) and quantified the number of ISA pixels within a 100-m-radius circular buffer around each nest box in QGIS (v 3.220; QGIS Development Team 2023) to represent territory size of this species during breeding (Wilkin et al. 2006). We computed the proportion of ISA by dividing the number of ISA pixels in this buffer by the total number of pixels (range: 0 – 1, where 1 = all ISA). We averaged the proportion ISA across nest boxes for a given study site to obtain a continuous urbanization metric at the site level (urban mean: 0.50, urban range: 0.21 – 0.98, forest mean: 0, forest range: 0 - 0).

### Common garden

During the spring of 2022, we conducted a common garden experiment by collecting eggs from the urban and forest great tit populations and rearing nestlings under standardized conditions (see also Thompson et al. 2024). We collected unincubated eggs that were covered and cold to the touch from urban and forest study sites between April 5 – 22 (N = 50 urban eggs from 4 sites, N = 40 forest eggs from 1 site). We collected between three and four eggs from each origin nest box (N = 23 origin nests) and transferred these eggs to wild foster nests at the Montpellier Zoo where females had just initiated incubation (N = 11 foster nests). The Montpellier Zoo is an established urban site with low to moderate levels of urbanization (average proportion ISA at 100m: 0.21) and is exposed to humans and associated stimuli. Foster nests contained eggs from two nests of origin (between six – eight eggs total) and, since urban origin lay dates tended to be earlier than forest ones (average 7.5 days earlier), we did not mix urban and forest eggs in the same foster broods. Of the 90 eggs transferred to foster nests, we had 73 nestlings hatch (N = 41 urban and 32 forest) that remained in foster nests until 10 days of age when they could thermoregulate on their own. We transferred 10-day old nestlings to captivity at the Zoo Nursery between April 29 – May 16 where they were ringed with a unique metal band and reared under the same captive conditions.

We initially reared nestlings in artificial nest boxes (i.e., open wooden boxes) that were kept in incubators to mimic a dark cavity (one to three broods per incubator). We fed nestlings every 30 minutes between 7:00 – 20:00 on a diet that consisted of hand-rearing powder solution (Nutribird A21 and A19, Versele-Laga, Deinze, Belgium), dead wax moth larvae, and mealworms. At 15 days old, nestling diet was enriched with a cake made of eggs, sunflower margarine, sugar, wheat and protein-rich pellet flours (Country’s Best Show1-2 Crumble, Versele-Laga, Deinze, Belgium) that was supplemented with commercial powders containing mostly vitamins and minerals (Nutribird A21, Versele-Laga; and Nekton-S, Nekton GmbH, Pforzheim, Germany). At approximately 18 days old, individuals began to “fledge” their captive nests and started flying around the nursery. We transferred these fledglings in the order they fledged to wire cages (2 – 3 individuals per cage irrespective of habitat of origin, foster brood, and sex) where we started to train individuals to feed by themselves. We initially fed individuals every 30 minutes, but at approximately 23 days old, feeding became less frequent (every hour, then every four hours) and individuals had access to food *ad libitum* in their cages. We considered birds independent at approximately 35 days old and in early June we transferred all individuals to large outdoor aviaries in randomly assigned groups of 6 – 8 birds blind to habitat of origin, foster brood, or sex. Birds were eating independently at this stage on a diet made of cake (see above) and live mealworms. Food and water were provided *ad libitum*. All common garden birds were reared by the same caretakers that were blind to habitat of origin as birds from urban and forest habitats were mixed throughout captive rearing. Temperature (range: 23 - 25°C) and humidity (60% or above) were kept constant through rearing.

### Cognitive assay

To measure and compare cognition in wild and common garden contexts we used the same cognitive assay. We designed a field assay similar to a motor detour task to evaluate inhibitory control and we administered this cognitive assay just before releasing individuals. Motor detour tasks have previously been adapted and administered successfully in great tits by, for example, using tasks with rewards not related to food or that do not require specific training steps (Isaksson et al. 2018; Davidson et al. 2022). One study showed that individual performance was repeatable across a modified detour task administered in the field and the classic detour “cylinder task” to measure inhibition in captivity (Davidson et al. 2022). We used a different field task that required individuals to escape a cage (i.e., the goal; Figure 1) by inhibiting their predominant impulse to fly into a transparent barrier in front of them and instead move laterally around the barrier to find the exit. We modeled the modified detour task after classic tasks used to measure inhibitory control but, unlike previous approaches, we did not have a habituation or training period. Thus, our task measures the ability of individuals to escape a challenging situation and their performance during the task likely reflects processes related to inhibitory control, exploration, and stress sensitivity.

**Fig. 1.**
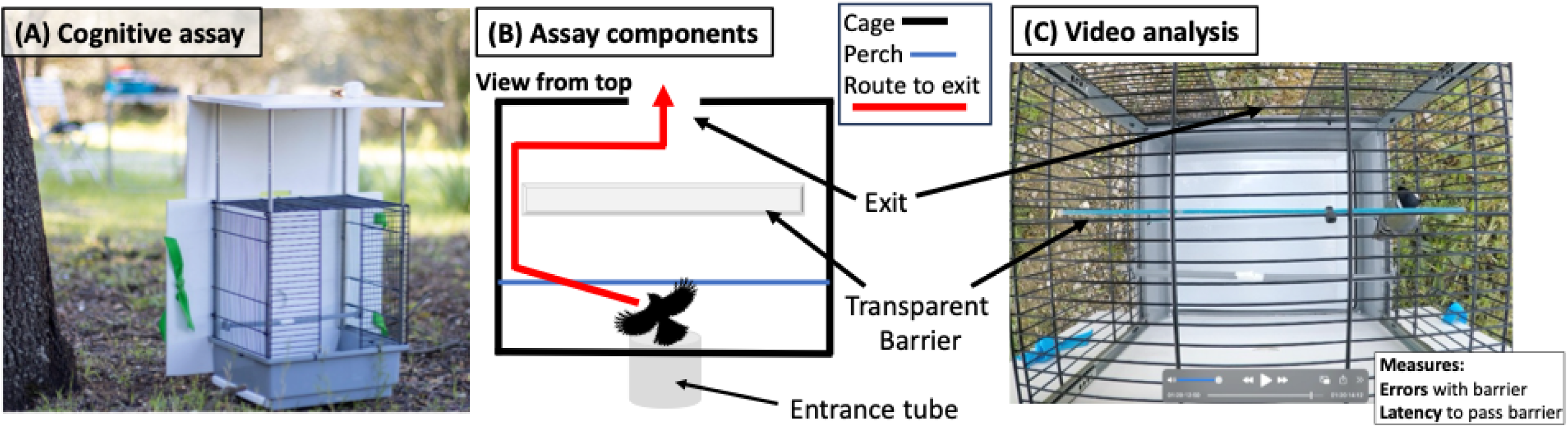
Cognitive assay used to measure performance related to inhibitory control in wild and common garden contexts. A) side view of the cognitive assay cage in the field, B) diagram showing assay components from above including a possible route to the exit in red, and C) a video clip showing how the cognitive task was analyzed including the measurements of interest (note bird detouring barrier on the right).

We first placed individuals in an enclosed opaque plastic tube on the outside of the cage to standardize their position before the assay and to give them one minute of standardized rest in the dark (Figure 1A). We then opened a sliding door that allowed access to the cage. In most cases, the bird entered the cage immediately and, in rare events where the bird did not enter, the observer would gently tap the end of the tube to encourage the individual into the cage. Upon entering, individuals were presented with an opening in the cage directly in front of them, but their direct path to this exit or goal was blocked by a transparent plexiglass barrier (Figure 1B). To escape the cage (i.e. the goal of the task), individuals needed to inhibit their predominant impulse to hit into the clear barrier and instead make a lateral motor detour towards gaps located at the side of the barrier to reach the exit. We allowed individuals up to 180 seconds to escape the cage before ending the trial and coaxing the bird to the exit.

We video recorded cognitive trials from above the cage (Figure 1C) that we later scored to extract two variables: the number of errors and the latency to escape. We recorded the number of errors as the number of hits an individual made into the transparent barrier during the trial before finding the exit (wild mean: 9.6 hits, wild range: 0 – 130; common garden mean: 4.9 hits, common garden range: 0 - 35). The average number of errors on the task was significantly lower in the common garden than in the wild (Welch’s t-test: t = 5.9, df = 553, P < 0.001). We also recorded the latency for individuals to complete the task as the amount of time in seconds from when >50% of the individual’s body entered the cage to the time when >50% of the body moved past the clear transparent barrier (wild mean: 22.81 sec, wild range: 0 – 180; common garden mean: 62.55 sec, common garden range: 0 - 180). The average latency to escape the cage was significantly higher in the common garden than in the wild (Welch’s t-test: t = -7.3, df = 255, P < 0.001). Videos were scored by two observers (DB: wild 2021; MJT: wild 2022 – 2023 and common garden) who had high inter-observer reliability (rho > 0.97) for both traits on a set of ten practice videos before initiating video analysis.

We compared individual performance between our field detour task and the classic cylinder task administered in captivity in a subset of individual great tits captured near Moulis, France (Crouchet 2024). The latency to escape, but not the number of errors, was significantly correlated between our field task and the classic cylinder task used to measure inhibition, suggesting that inhibitory control affected part of the response of the birds to our field assay (*N* = 15 individuals, latency: r^2^ = 0.52, *P* = 0.048; errors r^2^ = −0.19, *P* = 0.50).

In both wild and common garden contexts, we administered this assay after conducting a standardized phenotyping protocol where we measured other behavioural and morphological traits (Caizergues et al. 2018, 2022). In the wild, we conducted the assay in the vicinity of the nest box where individuals were captured. We placed the cage in a location facing away from roads or sidewalks in the wild, and the cage was positioned with the exit opposite to the sun to avoid reflections on the plexiglass window that could affect the bird’s response. In the common garden, we captured individuals for phenotyping using mist nets inside the large aviaries, and we conducted the cognitive assay on release back to the aviary. To visually and physically separate the focal individual from congeners that had already been released in aviaries, we used a thin sheet above the assay in the aviary that allowed light to pass through (similar to tree cover) and shaded the cage from the sun. Of the total number of individuals assayed (N = 393 wild and 73 common garden), N = 14 wild and 26 common garden individuals (4 and 36%, respectively) did not complete the task in 180 seconds in their first trial. In the common garden, we assayed individuals repeatedly and this improved to N = 5 individuals (8%) by the third trial.

### Blood sampling and DNA extraction

We took blood samples from individuals to determine their relatedness and exact nest of origin. This was necessary as foster broods contained eggs from two origin nests from the same habitat type and, apart from knowing whether individuals came from urban or forest habitats, we did not know the exact origin nest or relatedness of individuals after they hatched. We took blood samples a day before individuals were transferred to outdoor aviaries and we extracted DNA from these samples using Qiagen DNeasy blood and tissue kits. DNA extracts were sent to the Montpellier GenomiX platform (MGX) for RAD sequencing following the protocol described in Caizergues et al. (2022). We included N = 343 individual samples for sequencing, including 270 and 73 samples for wild and common garden individuals, respectively. Paired-end RAD- sequencing (2*150pb) generated 10.1 M reads with an average depth of 19.8x per individual before filtering. We used *STACKS v2.64* (Rochette et al. 2019) with the great tit genome reference (Laine et al. 2016) to call Single Nucleotide Polymorphisms (SNPs). Specifically, we ran *gstacks* with default options, except for --min_mapq=20, --rm-pcr-duplicates, and *populations* with default options except for --r=0.9, --max_obs_het=0.65, and --p=2. This step yielded 270,675 SNPs for 343 individuals (dataset 1). Based on this initial dataset, we created two different SNP datasets: one for the quantitative genetic analyses and one for the GWAS analysis.

First, for the quantitative genomic analyses we kept 185321 SNPs on autosomal chromosomes (dataset 2) after quality filtering (minimum average depth = 8, maximum average depth = 50, maximum 10% missingness per individual, LD < 0.5). Based on this SNP dataset, we estimated the genomic relatedness between all pairs of individuals (*N* = 343) using the make-rel option of the PLINK software (Purcell et al. 2007). We ensured the consistency of SNP calling by estimating the relatedness coefficient for five individuals and their associated technical replicates (i.e. one individual replicated once on a different lane) and ensuring that relatedness coefficients were close to one. The relatedness coefficient values between each pair of replicated samples were high, specifically 0.98, 0.99, 0.95, 0.99, and 0.99. For the quantitative genetics, we retained only one individual per pair. Additionally, we used the *ASRgenomics* package (Gezan et al. 2022) to perform a quality check on the genomic relatedness matrix (GRM) obtained with PLINK. The range of diagonal values was from 0.84 to 1.24, while the range of off-diagonal values was from −0.031 to 0.64. The variance of off-diagonal relatedness was 0.00099. We evaluated our statistical power to estimate heritability using this GRM by running a model analogous to that employed by Perrier et al. (2018) for morphological traits in blue tits. The similarity in heritability values obtained in our analysis suggests that our GRM has sufficient power. To maximize the number of individuals available for the quantitative genetic analysis, we used *ASRgenomics* to produce a hybrid matrix. This hybrid matrix, created by applying a correction on relatedness, allows the combination of relatedness from the social pedigree (obtained from observations in the field) and genomic relatedness into a single relatedness matrix called the Hybrid matrix (Hmatrix; Legarra et al. 2009; Christensen et al. 2012).

Second, to perform a Genome-Wide Association Study (GWAS), we constructed a SNP dataset that included autosomal and sex chromosomes, ensuring that the dataset contained SNPs with high linkage disequilibrium (LD). From the initial dataset comprising 270,675 SNPs across 343 individuals (dataset 1), we retained 248,325 SNPs on 243 individuals (dataset 3) after keeping individuals with a cognitive phenotype, removing unplaced scaffolds and applying the following filters: minimum average depth of 8, maximum average depth of 50, and a maximum of 10% missingness per individual. All bioinformatic analyses were conducted using a workflow built with the MBB-Framework (Penaud et al. 2020).

Finally, we built one last SNP dataset to identify the nests of origin of common garden birds. Following (Huisman 2017), we randomly subsampled 600 independent SNPs (dataset 4) with a minor Allele Frequency (MAF) of 0.4 to reconstruct the genetic pedigree between all 343 birds with *Sequoia* package (Huisman 2017). We could determine the nest of origin of individuals and the corresponding proportion ISA of origin study sites (continuous urbanization metric, see section ‘Study system’) using genetic relatedness between individuals. In one foster brood, we could determine relatedness among individuals (i.e., whether they were siblings) but not their exact nest of origin because parents from both origin nests had not been sampled. For individuals sharing this foster brood, we averaged the proportion ISA between the two possible urban origin sites.

### Statistical approach

We evaluated differences between urban and forest tits in their cognitive performance (number of errors and latency to escape) using R *v4.4.1* (R Core Team 2024). In both wild and common garden contexts, we used Bayesian mixed-effect models fitted with the *brms v2.21.0* package (Bürkner 2017) using Stan software (Carpenter et al. 2017). We analyzed wild and common garden data in separate models as they required different model structures. To address aim 1, we present four main models where we examine the two traits of interest in both contexts (model 1-4 below). To address aim 2, we present two quantitative genetic animal models that build on the models presented for aim 1 by adding the Hybrid matrix (pedigree + GRM relatedness) allowing the estimation of additive genetic variation and thus, heritability. We also address aim 2 by fitting two Genome Wide Association Analyses (GWAS) that are equivalent to linear (mixed) models testing the association between SNPs and cognitive traits.

We used the default priors proposed by brms for non-animal and animal models. We ran 4 chains for 20000 iterations each using a warm-up of 10000 iterations and a thinning interval of 1. Thus, model estimates and highest posterior density intervals (HPDIs) used posterior distributions consisting of 40000 samples. All models had appropriate convergence with Rhat < 1.01 (Vehtari et al. 2021), effective sample sizes >1000 (Bürkner 2017), and inspection of model diagnostic plots (traces, posterior predictive checks) confirmed good model fit. We interpreted estimates as having a clear effect when the HDPI (or credible interval) did not cross zero. We also report the probability of direction (pdirection) as additional information used to evaluate the certainty in effects, where pdirection = 0.97 indicates that 97% of the posterior distribution is the same sign as the median.

### Wild errors (model 1)

We evaluated the number of errors for individuals that successfully escaped the task within the 3-minute period. The number of errors was right-skewed and over dispersed due to a few large error values in both contexts (Figure S1). We therefore analysed the number of errors during the task using a negative binomial error structure. In the wild model, we tested whether there were differences between urban and forest birds in the number of errors they made by fitting habitat type (forest vs. urban) as a fixed effect. We also included trial (1-3), sex (female vs. male), and age (adult vs. juvenile), and their interactions with habitat type in the full model to determine whether there were different impacts across trials, sexes, or ages depending on habitat type. We also accounted for the date of testing (in Julian days since Jan 1), the time of day (continuous format: minutes divided by 60), year (2021 – 2023), and whether birds had a blood sample taken before testing (yes vs. no). We included individual ID (V_I_) and study site ID (V_SITE_) as random effects since some individuals had multiple trials and individuals were grouped within sites. We compared three models to determine which interactions to retain in the final model: a model with all interactions (between habitat type and sex, age, and trial), a model with an interaction between habitat type and trial, and a model with no interacting terms. The model with the lowest LOOIC (i.e., leave-one-out information criterion; similar to AIC) out of these three models was used as the final model (see Table S1). We calculated residual variance for negative binomial models following established methods (Tempelman and Gianola 1999; Mair et al. 2015) and estimated repeatability (*R*) as:

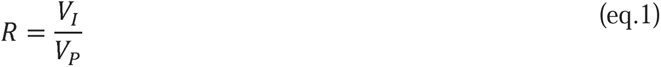

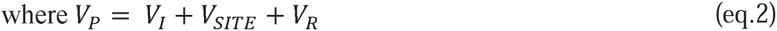

where V_I_ is the among-individual variance and V_P_ is the total phenotypic variance. V_P_ includes sources of among-individual variance (V_I_), among-site variance (V_SITE_), residual variance (V_R_).

We also ran three additional models: In the first model, we replaced habitat type with the proportion ISA to further evaluate how the number of errors may change with continuous urbanization (model structure determined via LOOIC; see Table S1). In the second model, we evaluated the robustness of our results by using only the first trial of the test across individuals since we had lower numbers of individual repeated measures in the wild. We used the same model structure and approach for this second model, but this model did not include trial as a fixed effect or individual ID as a random effect since there was only one trial per individual. In the third animal model, we added the Hmatrix as a random effect so we could estimate heritability (*h^2^*):

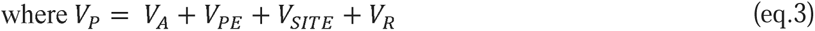

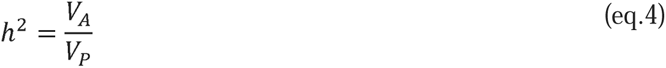

where V_A_ is the additive genetic variance and V_P_ is the total phenotypic variance. V_P_ comprises sources of additive genetic variance (V_A_; Hmatrix pedigree), permanent environmental variance (V_PE_; Individual ID), among-site variance (V_SITE_; site ID), and residual variance (V_R_).

### Common garden errors (model 2)

In the model run on the common garden birds, we also modeled the number of errors using a negative binomial error structure, but with a different model structure to account for the experimental design. We included habitat of origin as a fixed effect so we could determine whether birds from urban and forest habitats differed in the number of errors during the task. We included trial number (1-3) and sex (female vs. male) as fixed effects, and their interaction with habitat type to evaluate differential effects across habitat of origin. We also accounted for time of day (continuous format) as a fixed effect. We included the following random effects: nest of origin ID accounted for variance among origin nests (V_NO_), Individual ID accounted for variance among individuals (V_I_), foster nest ID accounted for variance among foster nests (V_NF_;), and aviary ID accounted for variance among social groups (V_AV_). Following the approach above (outlined for model 1), the final model retained interacting terms that achieved the lowest LOOIC (see Table S1). In an additional model, we substituted the habitat type effect with the proportion ISA. Repeatability of common garden models (*R_CG_*) was generated to account for a different random effect structure as follows:

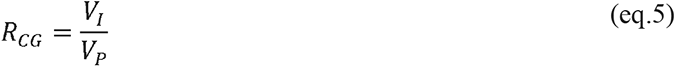

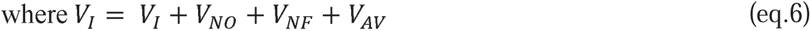

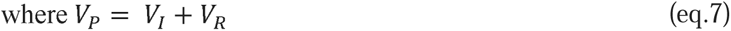

### Wild latency to escape (model 3)

The latency to escape the task was also right skewed with most individuals taking less than 50 seconds to escape and a few taking more time (Figure S1). As most wild individuals escaped the task (96%), we retained only successful individuals that escaped and modeled the latency to escape using a truncated lognormal error structure where the model was truncated at 180 seconds. We fit the same fixed and random effects as those included in our wild errors model (model 1). We followed the same approach as described for model 1 above: we evaluated the inclusion of interaction terms in the final model by identifying the model with the lowest LOOIC, we fit the same three additional models (ISA instead of habitat, trial one only, and an animal model with Hmatrix), and we calculated repeatability (using eqs.1-2) and heritability (using eqs.3-4) in the same way.

### Common garden latency (model 4)

Since some individuals did not escape the task in the common garden, there was a small peak in observations on the right extreme of the distribution (Figure S1). We excluded birds that did not escape the task (22% of observations) and modeled latency to escape the task as the response variable using a lognormal model truncated at 180 seconds. We included the same fixed and random effect structures used to evaluate the number of errors in the common garden (model 2). We also followed a similar model selection approach where interactions from the model with the lowest LOOIC was retained. Using the final model, we replaced the effect of habitat type with ISA in an additional model to evaluate how latency changed along the urban gradient. We generated repeatability of latency to escape in the common garden using eqs.5-7.

### Genome wide association study

To determine whether cognitive abilities are associated with SNPs and genes, we conducted a Genome Wide Association Study (GWAS) using the *v3.1.0 GAPIT* package (Lipka et al. 2012). First, to ensure that the phenotype data was normally distributed, an essential requirement in GWAS to avoid false positive results, we applied the rank-based inverse normal transformation, as recommended by Goh and Yap (2009) using the *RNOmni* package. We then analyzed a dataset containing 248325 SNPs and applied *GAPIT*’s Minor Allele Frequency (MAF) filter of 0.05, leaving 247940 SNPs for association testing in a sample of 253 wild individuals (Figure S2). *GAPIT* is designed to identify associations between SNPs and phenotypes across the genome by implementing several statistical models. Specifically, we employed several models provided by *GAPIT*, including the General Linear Model (GLM), Mixed Linear Models (MLM), which account for population structure and kinship, and the Bayesian Information and Linkage Disequilibrium Iteratively Nested Keyway (BLINK), which allows multi-locus tests. We assessed the fit of these statistical models by visually inspecting the QQ plots (Figure S3). We present only the results from the BLINK model as it demonstrated the best QQ plot fit and is known for having the highest power and accuracy among the statistical methods tested (Wang and Zhang 2021). Given that GWAS involves testing a large number of SNPs simultaneously, we applied the Benjamini-Hochberg procedure (i.e., False Discovery Rate method) to adjust p-values for multiple testing (Benjamini and Hochberg 1995). Finally, we identified whether the SNPs were within or near genes by selecting a 5kb window, based on the observation that in great tits, linkage disequilibrium falls to near baseline levels within 10kbp or less (Spurgin et al. 2024). To identify gene functions, we checked whether these genes were orthologous to others in the NCBI database, as orthologous genes typically have similar structures and, therefore, similar functions. We used the gene ontology provided by RefSeq to report gene functions and processes. Finally, to assess whether the significant SNPs explained a large portion of variation in inhibitory control, we fitted the same animal model as previously described in section ‘Wild errors (model 1)’, but with the significant SNPs coded as 0/1/2 (i.e., homozygous alternative allele, heterozygous and homozygous reference allele genotypes) as the only numeric fixed effects. We calculated the relative percentage of each variance component as previously, with the exception that we accounted for phenotypic variance due to SNPs in the denominator.

## Results

### Wild errors (model 1)

In the wild, urban and forest birds did not clearly differ in the number of errors made during the task (Figure 2A; CI crosses zero in Table 1.1A), but there was a tendency for urban birds to make fewer errors than forest birds (pdirection = 0.84, i.e., 84% of posterior distribution was positive). We found similar results when habitat type was replaced with continuous urbanization, where there was a tendency for birds to make fewer errors with increasing proportion ISA (pdirection = 0.91, Table S2.1A, but note an ISA*trial effect). The number of errors did not clearly differ across trials (Table 1.1A, but note pdirection = 0.82 for trial 3) and the above results were qualitatively similar when modelling the number of errors made during the first trial only (Table S3). The number of errors made by individuals was moderately repeatable (R = 0.38, CI = 0.23 – 0.53; Table 1.1A), suggesting that individuals consistently differed in the numbers of errors they made over repeated trials. Results were qualitatively similar to above when evaluating the number of errors in the wild using a quantitative genetic animal model (Table S4A). Although credible intervals were wide, the number of errors in the wild was moderately heritable (*h^2^* = 0.28, CI = 0.00 – 0.45, Figure 3A; Table S4A), while permanent environment (i.e., individual ID) and study site effects explained less variance in the number of errors (7 and 2% respectively, Figure 3A, Table S4A).

**Fig. 2.**
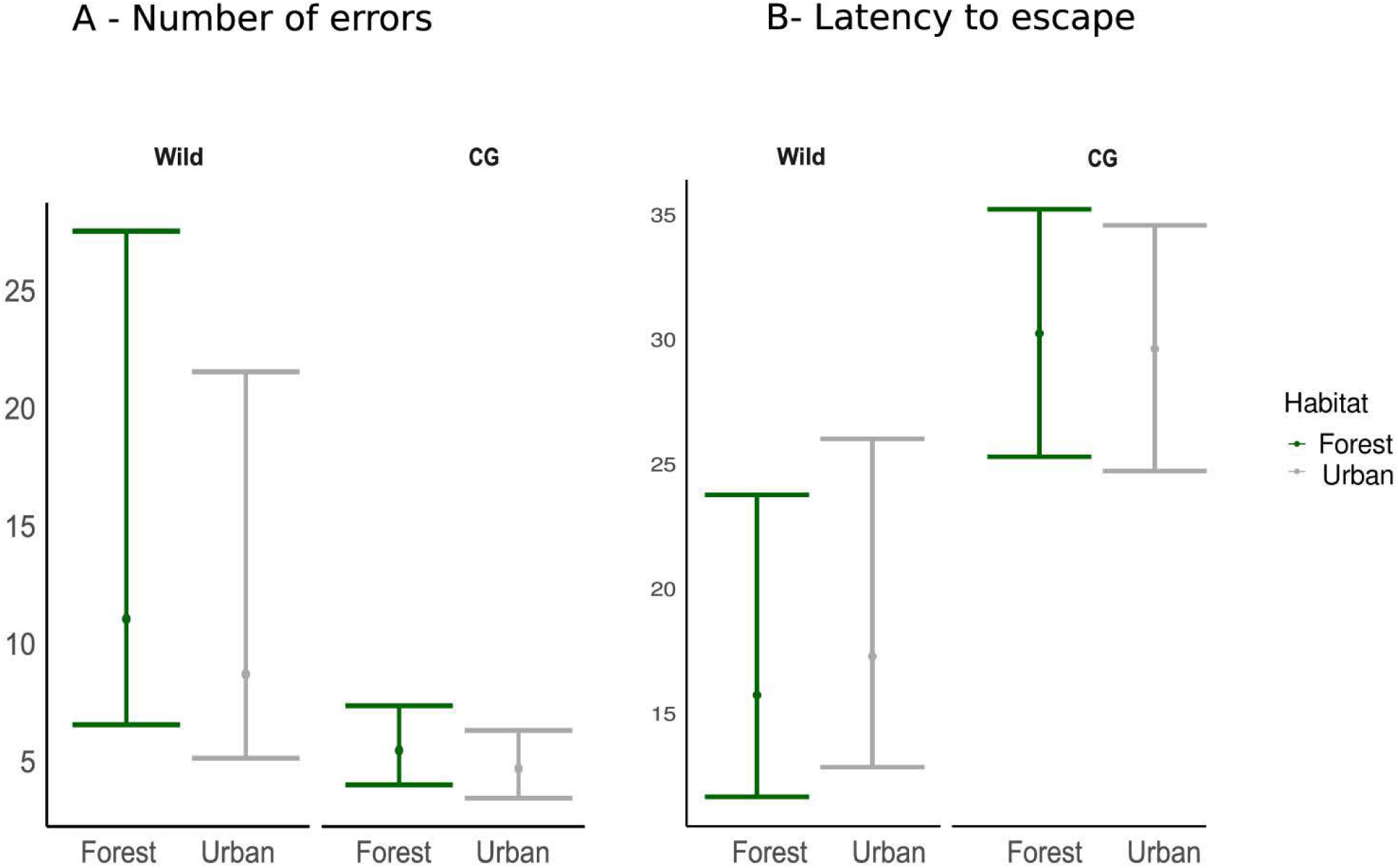
The effect of habitat type (forest vs. urban) and associated 95% credible intervals on the number of errors in wild and common garden (CG) contexts. Forest and urban estimates are shown in green and grey, respectively.

**Fig. 3.**
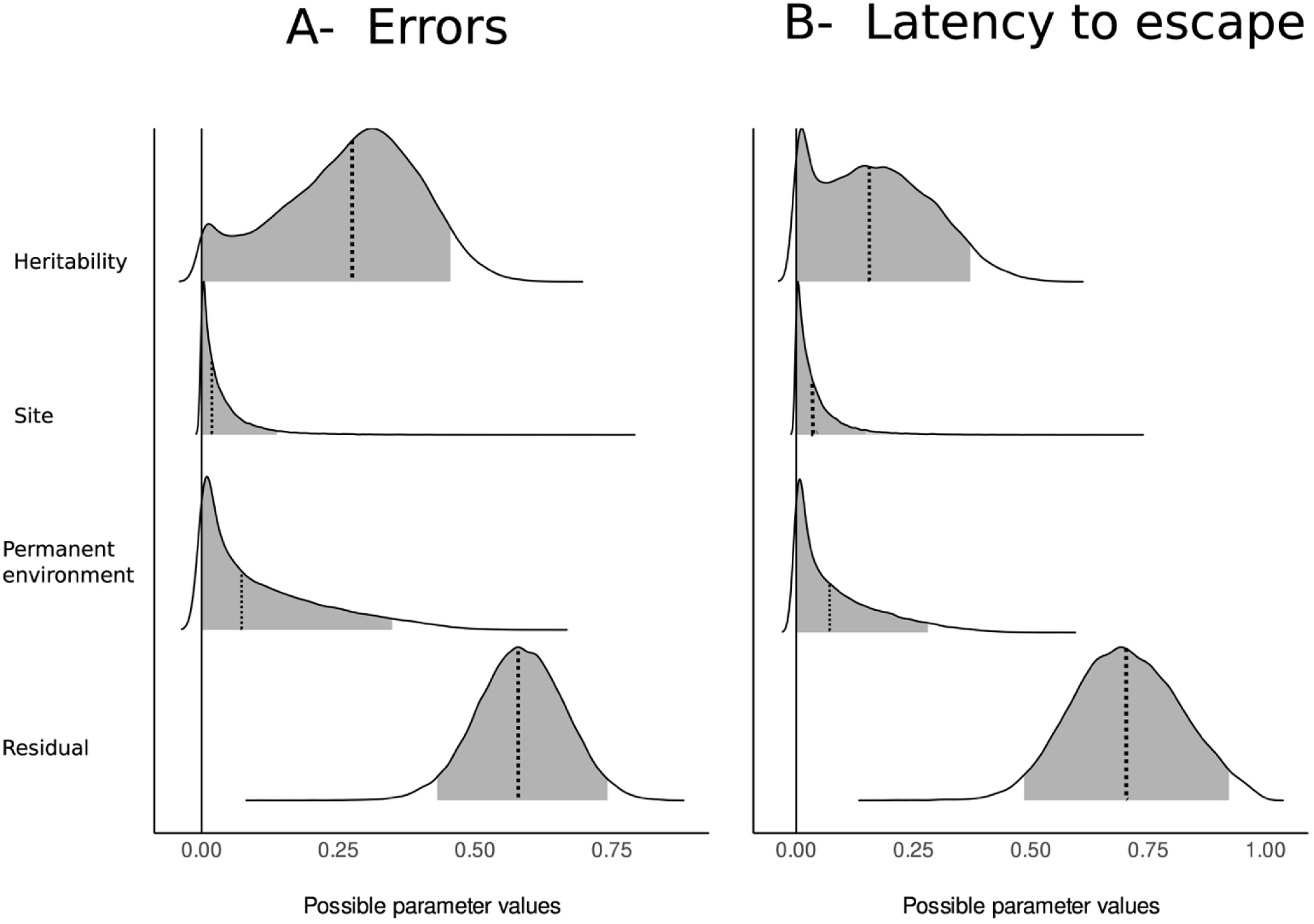
Highest Density Interval (HDI) of the posterior distributions of the ratio variance for errors (A) and latency to escape (B). The grey area represents the 95% HDI. From top to bottom, the posterior distributions are shown for heritability, site, permanent environment, and residual ratio variance. The solid line corresponds to zero. The dashed bold line corresponds to the median value.

**Table 1.** Posterior median and mean, credible intervals (CI or HDPI), and pdirection (probability effect is greater or less than zero) for fixed and random effects from separate 1) wild and 2) common garden contexts evaluating the A) number of errors and B) latency to escape in a motor detour task. The number of errors was fit with negative binomial generalized linear mixed-effect models and the latency to escape was fit with a truncated lognormal mixed-effect models. The number of observations (obs), individuals (ind), and individual repeated individual measures are shown for each context and trait. Fixed- and random- effect estimates whose credible intervals do not cross 0 or are greater than 0.001, respectively, are bolded.

| **1) WILD** | **A) ERRORS:** N = 442 obs, 380 ind<br>(56 ind – 2 trials, 10 ind – 3 trials) |  |  |  |  | **B) LATENCY:** N = 439 obs, 377 ind<br>(56 ind – 2 trials, 10 ind – 3 trials) |  |  |  |  |
| --- | --- | --- | --- | --- | --- | --- | --- | --- | --- | --- |
| *Fixed effects* | Median | Mean | CI |  | pdirection | Median | Mean | CI |  | pdirection |
| Intercept | **1.91** | **1.91** | **0.05** | **3.75** | **0.98** | **2.41** | **2.40** | **0.20** | **4.60** | **0.98** |
| Habitat (urban) | -0.21 | -0.21 | -0.75 | 0.33 | 0.84 | 0.14 | 0.14 | -0.53 | 0.82 | 0.72 |
| Sex (male) | 0.10 | 0.10 | -0.14 | 0.33 | 0.79 | 0.09 | 0.09 | -0.17 | 0.34 | 0.75 |
| Age (juvenile) | 0.06 | 0.06 | -0.19 | 0.31 | 0.69 | 0.08 | 0.08 | -0.19 | 0.36 | 0.71 |
| Julian date | 0.01 | 0.01 | -0.01 | 0.02 | 0.84 | 0.00 | 0.00 | -0.01 | 0.01 | 0.56 |
| Time of day | -0.03 | -0.03 | -0.10 | 0.03 | 0.83 | -0.03 | -0.03 | -0.11 | 0.04 | 0.83 |
| Year (2022) | -0.10 | -0.10 | -0.35 | 0.15 | 0.78 | -0.09 | -0.09 | -0.37 | 0.19 | 0.73 |
| Year (2023) | -0.10 | -0.10 | -0.40 | 0.21 | 0.73 | -0.02 | -0.02 | -0.35 | 0.33 | 0.54 |
| Blood (yes) | -0.27 | -0.28 | -0.62 | 0.06 | 0.95 | -0.23 | -0.23 | -0.60 | 0.16 | 0.88 |
| Trial (2) | 0.21 | 0.22 | -0.13 | 0.56 | 0.89 | 0.11 | 0.11 | -0.29 | 0.50 | 0.70 |
| Trial (3) | -0.34 | -0.34 | -1.06 | 0.41 | 0.82 | -0.56 | -0.56 | -1.38 | 0.26 | 0.91 |
| *Random effects* |  |  |  |  |  |  |  |  |  |  |
| Individual ID ($V_I$) | **0.47** | **0.47** | **0.30** | **0.67** | | 0.28 | 0.29 | 0.00 | 0.62 | |
| Site ID ($V_{SITE}$) | 0.02 | 0.05 | 0.00 | 0.17 | | 0.04 | 0.08 | 0.00 | 0.27 | |
| Residual ($V_R$) | **0.73** | **0.74** | **0.51** | **1.00** | | **1.22** | **1.22** | **0.84** | **1.60** | |
| Repeatability ($R$) | **0.38** | **0.38** | **0.23** | **0.53** | | 0.18 | 0.18 | 0.00 | 0.38 | |
| **2) COMMON GARDEN** | **A) ERRORS:** N = 153 obs, 72 ind<br>(54 ind – 2 trials, 32 ind – 3 trials) |  |  |  |  | **B) LATENCY:** N = 153 obs, 72 ind<br>(54 ind – 2 trials, 32 ind – 3 trials) |  |  |  |  |
| *Fixed effects* | Median | Mean | CI |  | pdirection | Median | Mean | CI |  | pdirection |
| Intercept | 1.06 | 1.05 | -0.86 | 2.93 | 0.86 | 2.99 | 3.01 | -0.40 | 6.81 | 0.95 |
| Habitat (urban) | -0.14 | -0.14 | -0.79 | 0.49 | 0.68 | -0.02 | -0.04 | -0.94 | 0.92 | 0.53 |
| Sex (male) | 0.20 | 0.20 | -0.24 | 0.62 | 0.82 | 0.31 | 0.31 | -0.43 | 1.00 | 0.81 |
| Time of day | 0.06 | 0.06 | -0.12 | 0.25 | 0.74 | 0.00 | 0.00 | -0.35 | 0.35 | 0.50 |
| Trial (2) | -0.28 | -0.28 | -0.74 | 0.15 | 0.90 | 0.13 | 0.14 | -0.69 | 1.00 | 0.62 |
| Trial (3) | **-0.42** | **-0.43** | **-0.88** | **0.01** | **0.97** | **-1.42** | **-1.43** | **-2.26** | **-0.57** | **1.00** |
| *Random effects* |  |  |  |  |  |  |  |  |  |  |
| Individual ID ($V_I$) | 0.22 | 0.24 | 0.00 | 0.52 | | 0.13 | 0.21 | 0.00 | 0.71 | |
| Origin nest ID ($V_{NO}$) | 0.05 | 0.08 | 0.00 | 0.29 | | 0.04 | 0.09 | 0.00 | 0.36 | |
| Foster nest ID ($V_{NF}$) | 0.03 | 0.08 | 0.00 | 0.30 | | 0.05 | 0.12 | 0.00 | 0.50 | |
| Aviary ID ($V_{AV}$) | 0.11 | 0.18 | 0.00 | 0.58 | | 0.29 | 0.45 | 0.00 | 1.47 | |
| Residual ($V_R$) | **0.89** | **0.92** | **0.51** | **1.44** | | **2.66** | **2.75** | **1.78** | **3.93** | |
| Repeatability ($R$) | **0.35** | **0.37** | **0.06** | **0.69** | | **0.22** | **0.24** | **0.02** | **0.51** | |

### Common garden errors (model 2)

In the common garden, the number of errors during the task was not affected by the origin habitat or level of urbanization (i.e., habitat or ISA CIs cross zero and pdirection < 0.75; Figure 2A; Table 1.2A; Table S2.2A). The number of errors clearly decreased with trial (Table 1.2A) where individuals made fewer errors during their last trial. Common garden birds were similarly repeatable (R = 0.35, CI = 0.06 – 0.69; Table 1.1A, see also Table S2.2A) to wild birds. Variance among individuals explained 15% of the variation in the number of errors, while differences between origin nests (3%), foster nests (2%), and aviaries (8%) explained less variation.

### Wild latency to escape (model 3)

In the wild, there were no clear differences in the latency to escape the detour task between urban and forest birds (Figure 2B; Table 1.1B) and no clear change over ISA (Table S2.1B; CIs cross zero and pdirection < 0.75). The latency to escape did not clearly change over repeated trials (Table 1.1B, but note pdirection = 0.91 for trial 3) and results were qualitatively similar when using data from only the first trial of individuals (Table S3B). The latency to escape the detour task was repeatable (R = 0.18, CI = 0.00 – 0.38; Table 1.1A), but less repeatable than the number of errors in the wild. Using the animal model, we estimated low heritability in the latency to escape (*h^2^* = 0.16, CI = 0.00 – 0.37; Figure 3B; Table S4B) relative to the number of errors, and permanent environment (i.e., Individual ID) and site effects explained little variation in this trait (<7%; Table S4B).

### Common garden latency to escape (model 4)

In the common garden, the latency to escape was not clearly affected by the origin habitat type or urbanization level (i.e., habitat or ISA CIs cross zero and pdirection < 0.75; Figure 2B; Table 1.2B; Table S2.2B). There was a clear effect of trial (Table 1.2B; see also Table S2.2B) where the latency to escape decreased clearly in the last common garden trial. Of the random effects tested, variance among individuals and aviaries explained the most variance in the latency to escape (4 % and 9%, respectively), while origin and foster nests explained negligible variation in this trait (1%, Table 1.2B). The latency to escape the detour task in the common garden was moderately repeatable (R = 0.22, CI = 0.02 – 0.51; Table 1.2B; see also Table S2.2B).

### Genome Wide Association

The GWAS revealed five SNPs and five genes significantly associated with the number of errors made during the task (Table 2, Figure 4A), but no SNPs were significantly associated with the latency to escape (Figure 4B). Significant SNPs associated with the number of errors were located on chromosomes 8, 13, 14, 23, and 24 collectively explaining 21% of the phenotypic variance (median = 0.21, CI = 0.12–0.29). However, the hybrid matrix explained an additional 7% of the phenotypic variance after accounting for variance explained by the five SNPs (i.e., the remaining genetic variance not explained by the SNPs or *h^2^_nonSNP =* 0.07); Table S5). The significant SNP on chromosome 14, explaining 4% of phenotypic variance, was within the Synaptogyrin 3 (SYNGR3) gene and showed a negative association with inhibitory control (effect size = −0.32). On chromosome 24, the significant SNP, explaining 4% of phenotypic variance was near two genes within a 5 kb window: Histone H4 Transcription Factor (HINFP) and POU Class 2 Homeobox-Associating Factor 2 (POU2AF2). The significant SNP on chromosome 13, explaining 7% of phenotypic variance, was also near two genes within a 5 kb window, though these genes do not have known functions (Table 2). Finally, the two remaining SNPs, located on chromosomes 8 and 23, explaining 2% of phenotypic variance each, were not linked to any known genes in the great tit genome. The functions of the identified genes and their associated processes are detailed in Table 2. The correlations between genotypes and the number of errors for each of the five SNPs are provided in the supplementary materials (Figure S4).

**Fig. 4.**
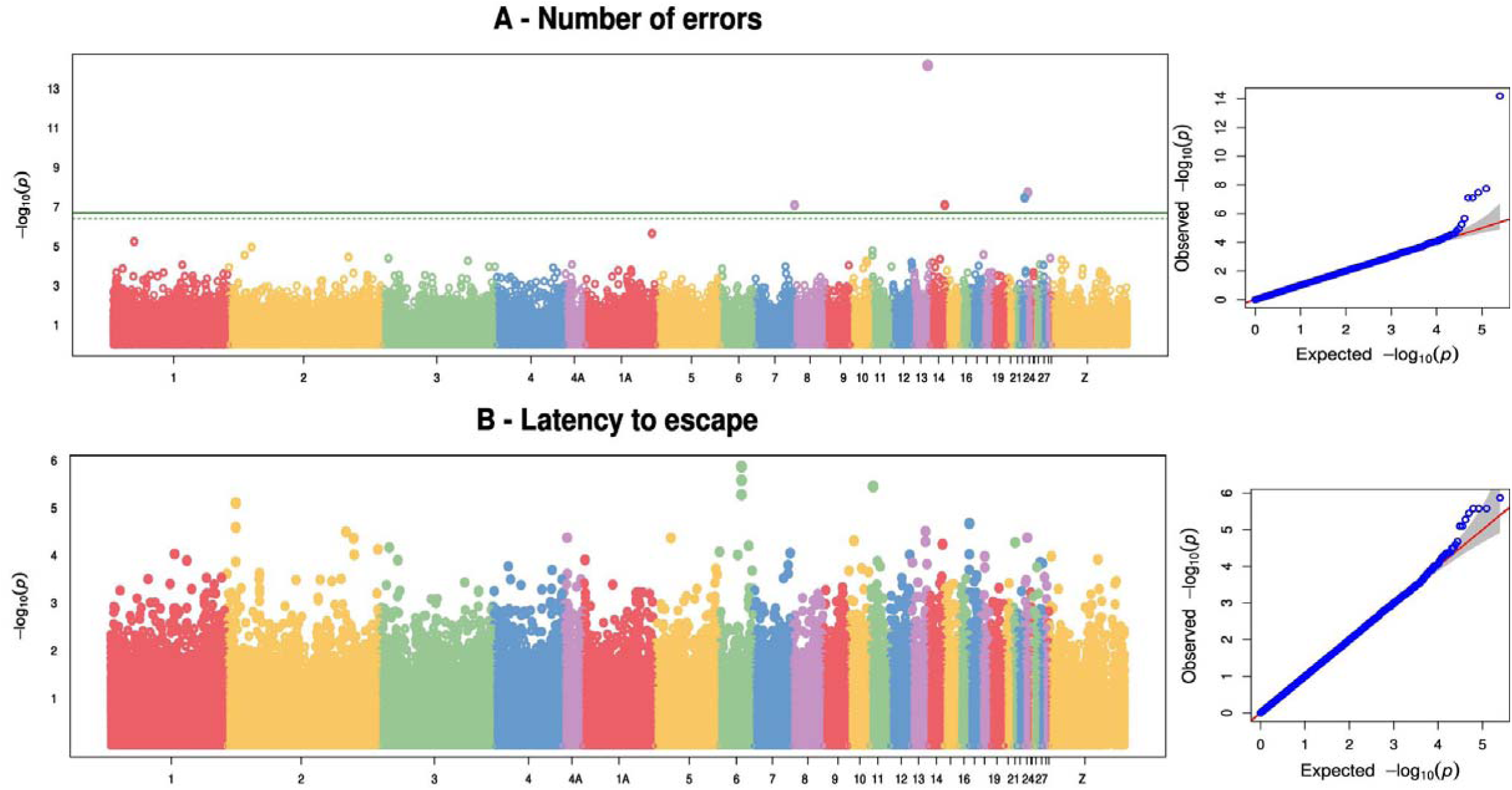
Manhattan plot (on the left) with –log10 p-values for the association between marker genotype and (A) number of errors, (B) Latency of escaping for all 228444 SNPs and 253 individuals and their associated Q-Qplot (on the right). On the left: The X-axis is the genomic position of the SNPs in the genome, and the Y-axis is the negative log base 10 of the P-values. The green solid line indicates the genome-wide Bonferroni-corrected significant threshold and the green dotted line the threshold for suggestive significant associations. Each chromosome is colored differently. SNPs with stronger associations with the trait will have a larger Y-coordinate value. On the right : Quantile-quantile (QQ) –plot of P-values. The Y-axis is the observed negative base 10 logarithm of the P-values, and the X-axis is the expected observed negative base 10 logarithm of the P-values under the assumption that the P-values follow a uniform [0,1] distribution. The grey surface show the 95% confidence interval for the QQ-plot under the null hypothesis of no association between the SNP and the trait.

**Table 2.** Single Nucleotide Polymorphisms (SNPs) and their associated genes identified by genome-wide analysis that were significantly associated with the number of errors. Information on chromosome number, position (in SNP base pairs), uncorrected *P- value* indicating the significance of the association between SNPs and the number of errors, the minor allele frequency of SNPs, number of genotypes, corrected H&B *P-value* (for multiple testing), and gene name (gene associated with the SNPs within a 5kb window). The sign of the allelic effect estimate is with respect to the nucleotide that is second in alphabetical order. Note: There were no significant SNPs associated with latency to escape.

| Trait | Chromosome | Position | *P-value* | Minor allele frequency | Allele Frequencies | # genotypes | H&B *P-value* | Alleles | Effects | Gene name | Function | Process |
| --- | --- | --- | --- | --- | --- | --- | --- | --- | --- | --- | --- | --- |
| Errors | 13 | 14568042 | 0.000 | 0.23 | CITY: 0.240<br>FOREST: 0.196 | 253 | 0.000 | C/T | 0.59 | LOC107210618<br>LOC117245099 | Unknown | Unknown |
| Errors | 24 | 1757705 | 0.000 | 0.48 | CITY: 0.410<br>FOREST: 0.588 | 253 | 0.002 | G/T | 0.33 | HINFP | -DNA-binding transcription factor activity, RNA polymerase II-specific<br>-RNA polymerase II cis-regulatory region sequence-specific DNA binding (Gbinding(allus gallus)) | -Anatomical structure development<br>-Regulation of transcription by RNA polymerase II<br>-Orthologous to HINFP in humans and mice |
| Errors | 24 | 1757705 | 0.000 | 0.48 | CITY: 0.410<br>FOREST: 0.588 | 253 | 0.002 | G/T | 0.33 | POU2AF2 | -POU domain binding enables protein binding, sequence-specific DNA binding, transcription coactivator activity (humans) | -positive regulation of DNA-templated transcription |
| Errors | 23 | 5329402 | 0.000 | 0.48 | CITY: 0.503<br>FOREST: 0.429 | 253 | 0.003 | C/G | -0.32 | NA |  |  |
| Errors | 8 | 675877 | 0.000 | 0.32 | CITY: 0.321<br>FOREST: 0.326 | 253 | 0.004 | A/G | -0.32 | NA |  |  |
| Errors | 14 | 14857470 | 0.000 | 0.11 | CITY: 0.101<br>FOREST: 0.134 | 253 | 0.004 | C/T | -0.47 | SYNGR3 | - SH2 domain binding and protein binding (humans) | -regulation of neurotransmitter uptake (IEA), substantia nigra development, regulated exocytosis, positive regulation of transporter activity |

## Discussion

We aimed to address two major unknowns in the literature about whether urban and nonurban individuals differ in inhibitory control on a detour task, and whether genetic variation underlies cognitive variation observed along an urbanization gradient. Despite growing interest in the evolutionary potential of cognitive traits and the relevance of inhibitory control for adjusting to urban conditions, no studies have yet examined this cognitive trait in wild urban populations. We found limited evidence that urbanization (categorical and continuous) was associated with performance related to inhibitory control during a detour task. This finding was echoed in the common garden experiment as birds from urban and nonurban origins did not clearly differ in their performance on the same task. We found that performance related to inhibitory control was moderately repeatable in the wild (*R* = 0.38 and 0.18 for errors and latency to escape, respectively) and some evidence for genetic variance in inhibitory control performance contributing towards low to moderate heritability (*h^2^* = 0.28 and 0.16 for number of errors and latency to escape, respectively). We also found that the ability to inhibit the number of errors during the task was significantly associated with five candidate genes explaining >20% of cognitive variation, further supporting that inhibitory control in the wild is partially genetically controlled. Overall, we find support that cognitive variation related to inhibitory control has a genetic basis in the wild, but that urban populations do not phenotypically or genetically differ from their forest counterparts in this cognitive trait.

Despite high heterogeneity among studies, urban animals tend to have higher performance on cognitive tasks (e.g., problem solving; Vincze and Kovács 2022). We did not find support for higher urban cognitive performance as urban and forest great tits did not clearly differ in the number of errors or latency to escape a motor detour task. In the predicted direction, there was a weak tendency (i.e., 91% of posterior distribution > 0) for tits in more urbanized habitats to make fewer errors than tits in less urbanized areas, but this trend was statistically weak (credible interval crosses zero; Table S2). It is therefore possible that urban great tits have higher inhibitory control performance than forest tits, but our results suggest that this difference is at best small (i.e., birds make 1.5 fewer errors on average in habitat with the highest ISA compared to the lowest ISA). Our assay may not exclusively measure inhibitory control, and other cognitive processes (and non-cognitive ones like motivation and experience) could impact the number of errors and the latency to escape the cage during our task. For example, a previous study showed that the number of errors made by individual pheasants during detour tasks may be more representative of individual measures of persistence (van Horik et al. 2018; see also Prasher et al. 2019), which may suggest that urban tits are marginally less motivated to escape than forest tits. We do not have information on the past experience of each individual but, as urban animals may be exposed to more non-natural surfaces like windows in their environments, experience with transparent barriers could play a role in how urban animals respond to detour tasks (van Horik et al. 2018; Isaksson et al. 2018). Prior comprehension of transparent barriers or higher persistence could thus explain why urban great tits made slightly fewer errors during the task than forest tits.

Although we did not find clear differences in inhibitory control between wild and forest great tits, countergradient variation can mask phenotypic divergences between wild populations (Conover et al. 2009). For instance, genetic change may drive higher cognitive performance in urban populations, while environmental conditions in urban areas could impair cognitive performance (e.g., lower quality food sources or noise pollution; Lee and Thornton 2021; Templeton et al. 2023), so that divergence in cognition between urban and nonurban populations are not detected. Using a common garden experiment allowed us to confirm that countergradient variation was not acting on inhibitory control in these wild great tits. Birds from urban and forest origins did not differ in their performance related to inhibitory control and, thus, we did not find evidence that urban and forest great tits differ genetically in inhibitory control. Cognitive traits like learning, problem solving, or inhibitory control can buffer wild populations from environmental changes and “buy time” for newly established populations to adapt to urban conditions (Caspi et al. 2022). However, urban great tits do not appear to clearly differ (phenotypically or genetically), on average, from their forest counterparts in their performance related to inhibitory control, so this trait may not play a large role in adaptation to urban environments (but see Thompson et al. 2024 for evidence of maintained differences in other trait types). Urbanization can also affect variation in traits with cascading ecological and evolutionary consequences (Alberti et al. 2017; Thompson et al. 2022), and so evaluating whether differences in individual variation between urban and nonurban habitats could further help to establish how urbanization and cognitive abilities are related.

Performance in the detour task differed across wild and common garden contexts where birds in the common garden on average took more time to escape the cage and made fewer errors than birds in the wild. Differences in motivation or stress between the wild and common garden could explain why performance differed between these contexts. Individuals in the common garden were reared for part of their life in cages and so common garden birds may be less motivated or stressed than wild birds to escape the detour task (i.e., the goal) because they are used to being caged. Less motivated individuals would have more time to evaluate the task and perceive the transparent barrier thereby increasing their time to escape while reducing the number of errors with the barrier before escaping. Another difference between contexts related to the improvement in performance over repeated trials of the task. We did not find clear evidence that performance in the wild improved over trials, but we did see significant declines in the number of errors and the latency to escape the cage in the common garden experiment. In the common garden, we assayed most individuals every three months, whereas in the wild we administered the task annually and had fewer repeated individual measures. Improved performance in the common garden could indicate that individuals learned the task or that their cognitive performance increased with age between 74 and 264 days old. In the wild, however, a lack of improvement could suggest that longer time periods between testing reduces learning or that motivation or stress levels during wild testing alter responses.

We found that individuals were consistently different among each other over time (i.e., repeatable) in the number of errors and latency to escape the task in the wild (*R =*0.38 and 0.18). Our estimates of repeatability for inhibitory control are slightly higher than repeatability estimates of cognitive traits reported in a meta-analysis (*R* = 0.15 – 0.28; Cauchoix et al. 2018) and were also higher than repeatability estimates of inhibitory control in another study using wild great tits (*R* = 0.15; Davidson et al. 2022). The latency to escape our task and the latency to a reward in a classic cylinder detour task were positively correlated in a different subset of birds, suggesting that our task is measuring processes related to inhibitory control, but in future we will need to validate our cognitive measures by comparing individual performance across different inhibitory control tasks (i.e., convergent validity; Völter et al. 2018; Davidson et al. 2022). Estimating repeatability is a useful first step in studies seeking to evaluate the evolutionary potential of traits as repeatability quantifies variance related to among-individual differences and can be used to infer the upper limit of heritability (Wilson 2018).

We were also able to estimate the heritability of inhibitory control traits in the wild and, in line with previous work, our estimates ranged from low (latency to escape *h^2^*= 0.16) to moderate (number of errors *h^2^* = 0.28). However, we lacked power (note bimodal posterior distribution in Figure 4, Pick et al. 2023) to confidently conclude that our heritability estimates were greater than zero, which is unsurprising given that the number of repeated individual observations we have in our data is low (Jablonszky and Garamszegi 2024). Estimating repeatability or heritability of cognitive traits in wild populations is challenging as the context of measurement can greatly influence individual responses and since experience or learning can affect performance in subsequent assays (Cauchoix et al. 2018, 2020). Most estimates of heritability for cognitive traits in the wild tend to be low and hardly distinguishable from zero, and most studies conclude that cognition is mainly environmentally determined (Van Den Heuvel et al. 2023; McCallum and Shaw 2023; Speechley et al. 2024). For example, in wild New Zealand toutouwai (North Island robins; *Petroica longipes*), a quantitative genetic study revealed that repeatable individual variation in inhibitory control had little genetic basis (McCallum and Shaw 2023). Although we lack certainty around our heritability estimates, we provide contrary evidence supported by genomic results that performance related to inhibitory control has a genetic basis in wild great tits. Determining how inhibitory control relates to fitness metrics in the wild (Morand-Ferron et al. 2016) would be an obvious next step to further evaluate whether this trait is selected for.

The serotonergic and dopaminergic systems, and their interaction, play a crucial role in shaping cognitive abilities, including motor control, learning, memory, and aversive or exploratory behaviours (Durstewitz et al. 1999). These systems are remarkably conserved across mammals and birds (Durstewitz et al. 1999; Bacqué-Cazenave et al. 2020; Fujita et al. 2023), and play a role in inhibitory control (Weafer et al. 2017). In our study, we found that inhibitory control in the great tit was also associated with genes involved in the serotonergic and dopaminergic systems, specifically the Histone H4 Transcription Factor (HINFP) and Synaptogyrin 3 (SYNGR3). These genes have been shown to play significant roles in cognitive functions in humans and mice and are implicated in Alzheimer’s disease (Emilsson et al. 2006; Saetre et al. 2011; Gupta and Kumar 2021). Interestingly, we did not find support for genes that have previously been identified as being associated with cognitive traits in non-human organisms. For example, while three SNPs in the serotonin transporter gene (SERT) have been linked to problem-solving in great tits (Grunst et al. 2021), the SNPs identified in the SERT gene in our study were not significantly associated with performance related to inhibitory control. However, in line with Grunst et al. (2021), we found that a few SNPs and genes were linked to cognitive abilities, suggesting that inhibitory control might have an oligogenic basis. Note however that the five significant SNPs identified accounted for 21% (CI = 12 -29) of the variation in the number of errors, leaving only 7% of the genetic variance related to inhibition unexplained by the GWAS. This suggests that additional causal variants may exist, but their effects are either too small to meet stringent significance thresholds or are not in complete linkage disequilibrium with the genotyped SNPs (Yang et al. 2010). Recent studies on moderately heritable traits, such as morphological traits, have demonstrated the polygenic nature of these traits in wild populations, even in the absence of GWAS peaks. For instance, studies with large sample sizes in red deer (Gauzere et al. 2023; N > 2000 *Cervus elaphus*) and hihi (Duntsch et al. 2020 N > 500 *Notiomystis cincta*) were unable to identify significant genes using the classical GWAS approach, but demonstrated the polygenic nature of traits using other methods. It is therefore possible that different approaches could reveal that inhibitory control in great tits has a polygenic determinism in the future. Another study on cognitive abilities in only a few hundred mountain chickadees (*Poecile gambeli*) managed to show a polygenic basis for spatial memory using GWAS, identifying over 100 associated genes using whole-genome sequencing (Semenov et al. 2024). This is not surprising, as whole genome sequencing offers greater statistical power for detecting polygenic traits compared to RAD sequencing (Lowry et al. 2017), which, in our specific case, covered an average of only 4% of the genome. The reduction in costs for whole-genome sequencing presents exciting opportunities to further explore the genetic architecture of cognitive abilities moving forward. This approach could help determine whether a polygenic structure hinders or promotes evolutionary processes, shedding light on the complex genetic underpinnings of cognitive traits.

In conclusion, urban and forest great tits do not differ (phenotypically or genetically) in their cognitive abilities related to inhibitory control, but we did find evidence that inhibitory control has a genetic basis within these wild populations. Previous work highlights mixed results concerning how urbanization impacts cognitive abilities (Sol et al. 2020; Vincze and Kovács 2022). Cognitive expression is likely context dependent (Cauchoix et al. 2020) meaning that the association between urbanization and cognition will likely also depend on the cognitive trait, species, and environmental axis of urbanization studied. Identifying specific selection pressures that act on cognition, which may or may not differ between urban and nonurban settings (Charmantier et al. 2024), will be crucial for establishing the role of cognition in urban evolution. Although heritability of cognition is more difficult to measure and detect in wild than laboratory populations, there is accumulating evidence that cognition can be partially genetically determined in the wild. Together our results on the heritability and genomics of inhibitory control in great tits provide further support that cognitive performance has a genetic basis and, thus, has the potential to evolve in the wild. New and more affordable genomic methods provide exciting opportunities to explore the genetic architecture of cognitive traits in a variety of contexts and species. These approaches will have especially meaningful implications for evaluating the role of cognition in adaptation to novel contexts and the long-term persistence of populations in cities.

## Acknowledgments

We would like to thank everyone who participated in the common garden experiment including Barbara Class, Segoléne Delaitre, Amélie Fargevieille, Lisa Sandmeyer, Christophe de Franceschi from the CEFE and Baptiste Chenet, Marc Romans, Vivian Espinasse, Thibault Pujol, Flavien Daunis, Cathie Troussier, Jérôme Brière, Laetitia Boscardin, Sébastien Pouvreau, Lucas Boussioux, Charlotte Gay, Marion Darde from the Montpellier Zoo. We would also like to thank all the people that participated in the field monitoring in Montpellier and La Rouvière including Arnaud Grégoire, Marcel Lambrechts, Samuel Perret, and Paul Cuchot. Additional thanks to those in Moulis including Clea Lecorre, Renan Destrades, Thomas Crouchet, Philipp Heeb, Joel White and Thomas Deruelles. We would like to thank Benjamin Penaud who provided advice and helped with the bioinformatic workflows. MJT was supported by a Canadian Graduate Scholarship from the Natural Sciences and Engineering Research Council (NSERC) of Canada, a Fonds de recherche du Québec Nature et Technologies PhD scholarship, and a PhD mobility grant from le Centre Méditerranéen de l’Environnement et de la Biodiversité (CeMEB). This project was funded by the Agence Nationale de la Recherche (URBANTIT grant ANR-19-CE34- 0008-05 and SoCo grant ANR-18-CE02-0023 to AC and ASC), the GenCog project funded by the Laboratoire d’Excellence TULIP (ANR-10-LABX-41) to LG, ASC, CP and AC, and a Fonds de Recherche du Quebec Nature et Technologie to DR. The long term Montpellier monitoring is supported by the OSU-OREME and part of the long-term Studies in Ecology and Evolution (SEE-Life) program of the CNRS.

