## Supplementary materials for "Tit wit: environmental and genetic drivers of cognitive variation along an urbanization gradient"

**Title:**

*Joint first author

**Affiliations:**

**Corresponding Author:**

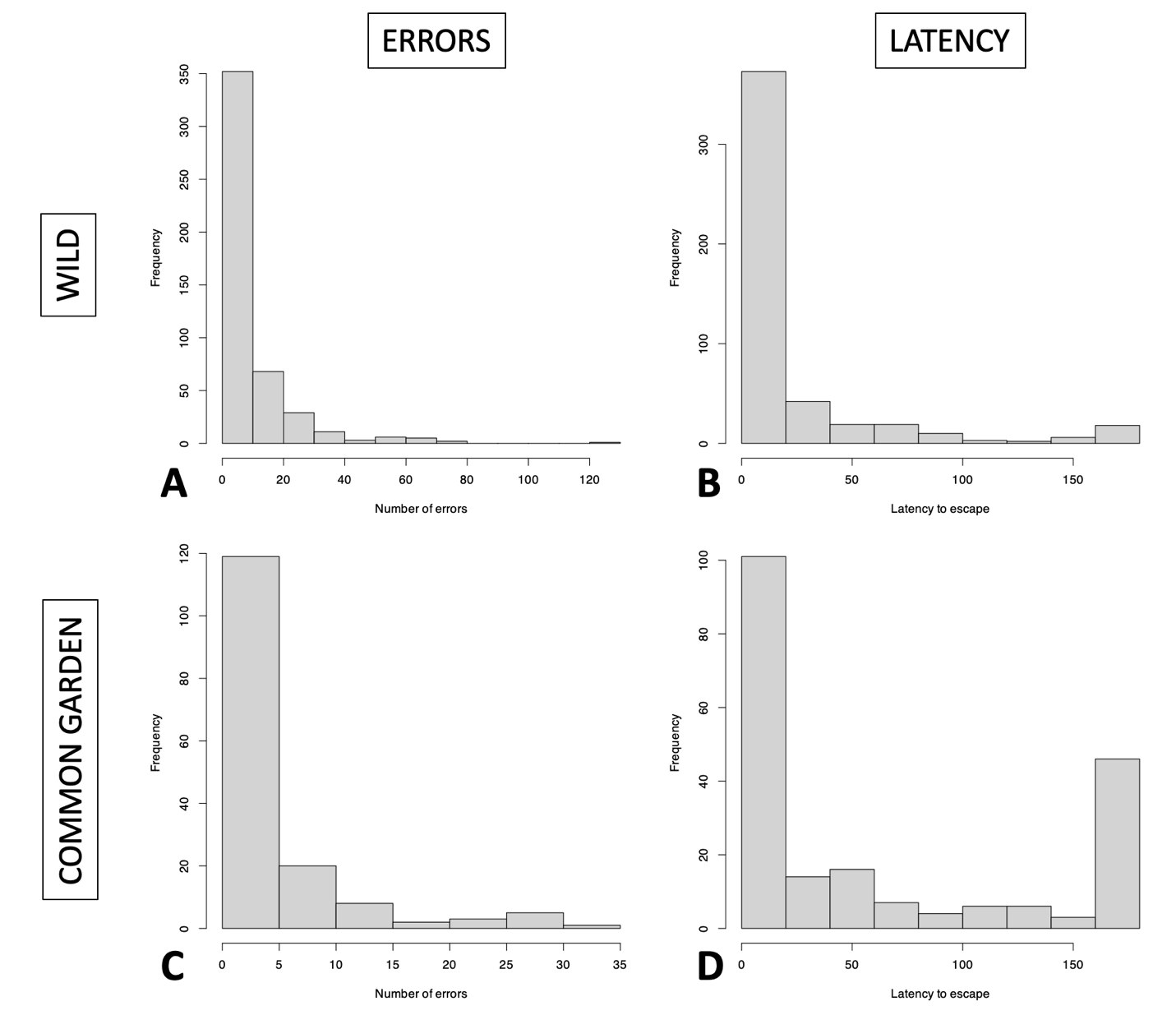

**Fig.S1** Distributions of the number of errors (A & C) and latency to escape the cage (B & D) in the motor detour task administered in wild (top panel) and common garden (bottom panel) contexts.

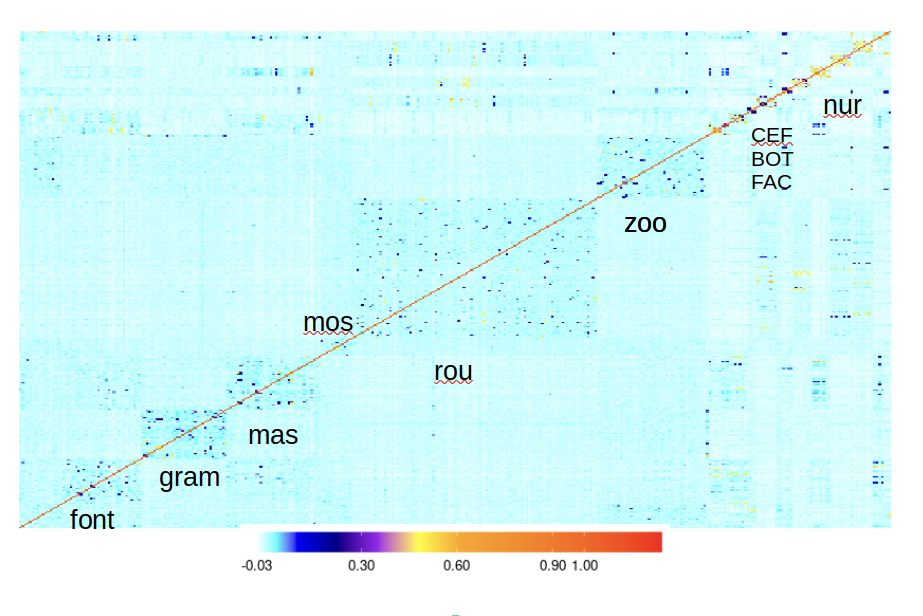

**Fig.S2** Genomic relatedness heatmap, with relatedness represented by a color gradient ranging from cold colors (blue with low relatedness) to hot colors (red with high relatedness). The names of study sites and contexts are labeled in black (urban: FONT, GRAM, MAS, MOS, ZOO, FAC, CEFE, BOT; forest: ROU; common garden birds: NUR).

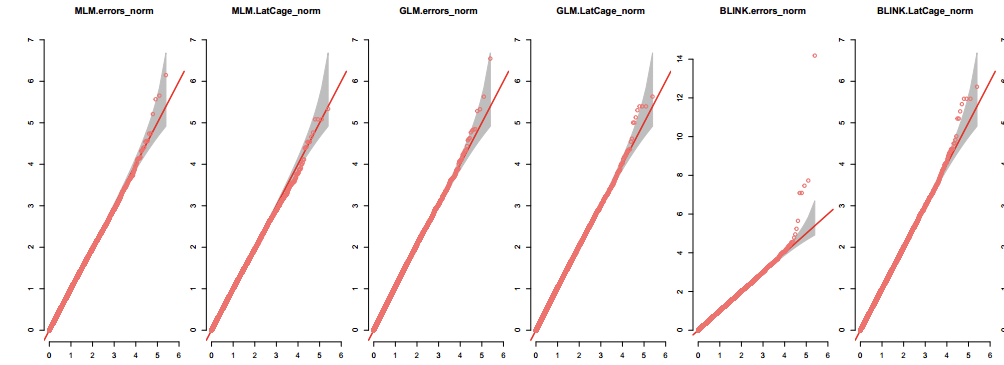

**Fig.S3** Quantile-quantile (QQ) plots of *P*-values for the different GWAS models: MLM, GLM and BLINK for number of errors and latency to escape the task. The y-axis is the observed negative base 10 logarithm of the *P*-values and the x-axis is the expected observed negative base 10 logarithm of the *P*-values under the assumption that the *P*-values follow a uniform [0,1] distribution. The grey surface shows the 95% confidence interval for the QQ plot under the null hypothesis of no association between the SNP and the trait.

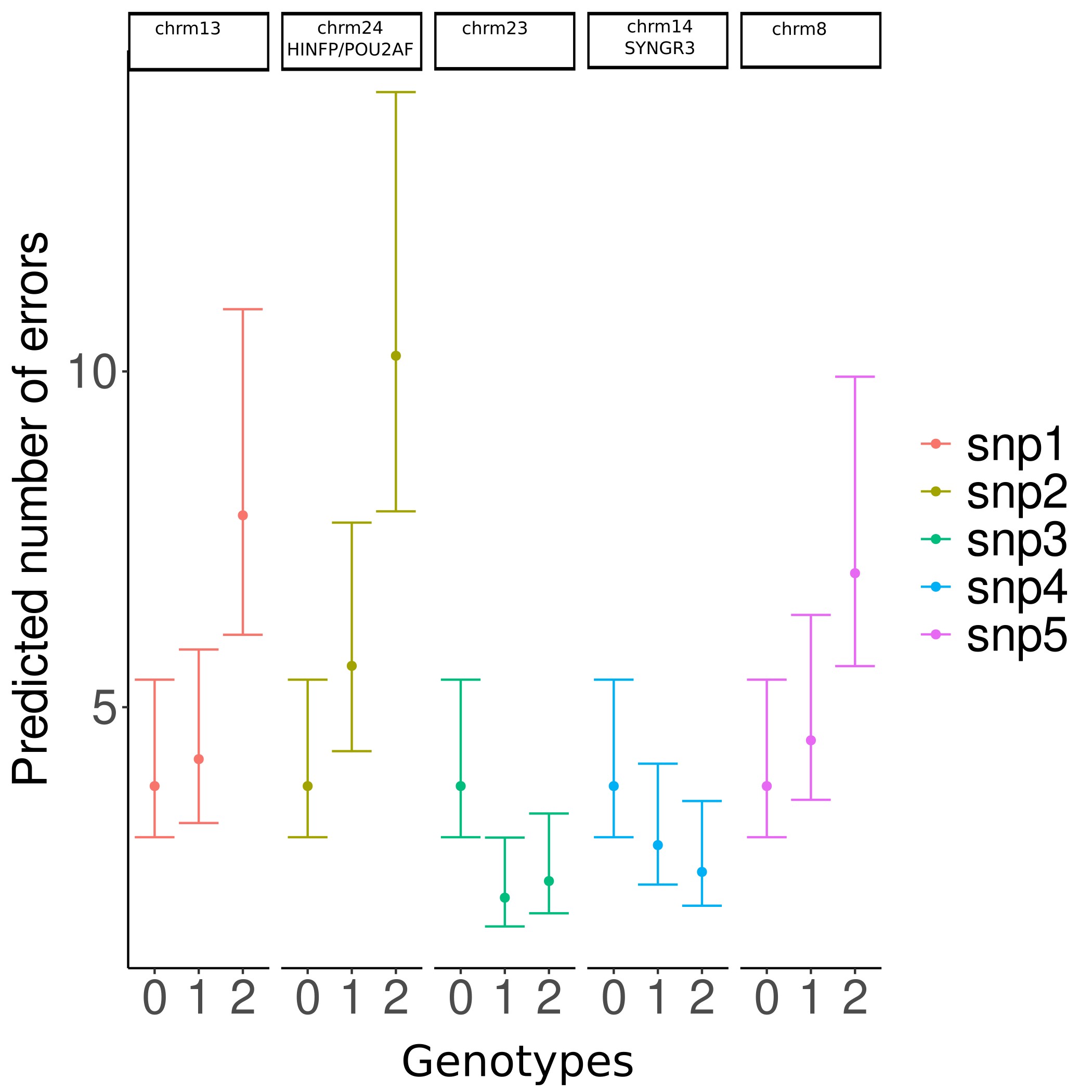

**Fig.S4** Relationship and 95% credible interval between genotypes and number of errors for significant SNPs. Genotypes are coded as 0, 1, and 2 for homozygous alternative alleles, heterozygotes, and homozygous reference alleles, respectively. Note: There were no significant SNPs associated with latency to escape.

**Table S1.** Leave-one-out cross validation information criterion (LOOIC) and their corresponding standard errors (in square brackets) from models comparing interaction term structures for both the A) number of errors and B) latency to escape in wild and common garden contexts (see main text where each model number is fully described). Instances where ISA* or habitat* are followed by (trial+sex+age) indicates that two-way interactions were fit with ISA or habitat and all these terms.

| Model structure: | A) ERRORS | B) LATENCY |
| --- | --- | --- |
| *Wild* | *Model 1 - ISA* | *Model 3 - ISA* |
| No interactions | 2871.21 [50.40] | **3257.94 [62.71]** |
| ISA*trial + others fixed effects | **2866.39 [50.19]** | 3259.05 [62.63] |
| ISA*(trial+sex+age) + others fixed effects | 2867.58 [49.88] | 3262.45 [62.41] |
| *Wild* | *Model 1 - habitat* | *Model 3 - habitat* |
| No interactions | **2867.30 [49.97]** | **3257.45 [62.59]** |
| Habitat*trial+ others fixed effects | 2870.49 [50.22] | 3261.41 [62.44] |
| Habitat*(trial+sex+age) + others fixed effects | 2874.07 [50.36] | 3264.12 [62.34] |
| *Common garden* | *Model 2 - ISA* | *Model 4 - ISA* |
| No interactions | **822.55 [31.22]** | **1269.70 [36.86]** |
| ISA ISA*trial+ others fixed effects | 824.78 [31.56] | 1274.65 [36.50] |
| ISA*(trial+sex+age) + others fixed effects | 826.33 [31.39] | 1276.14 [36.20] |
| *Common garden* | *Model 2 - habitat* | *Model 4 - habitat* |
| No interactions | **823.70 [31.37]** | **1269.66 [36.94]** |
| Habitat*trial+ others fixed effects | 823.88 [31.43] | 1273.33 [36.42] |
| Habitat*(trial+sex) + others fixed effects | 826.42 [31.66] | 1274.85 [36.32] |

**Table S2.** Model comparison to Table 1 in main text when examining the site-level proportion ISA effect (impervious surface area; 100m) instead of the habitat type (forest vs. urban) effect. Posterior median and mean, credible intervals (CI), and pdirection (probability effect is in the same direction as median) for fixed and random effects from separate 1) wild and 2) common garden contexts evaluating the A) number of errors and B) latency to escape in a motor detour task. The number of errors was fit with negative binomial generalized linear mixed-effect models and the latency to escape was fit with a truncated lognormal mixed-effect models. The number of observations (obs), individuals (ind), and individual repeated individual measures are shown for each context and trait. Estimates whose credible intervals do not cross 0 are bolded.

| **1) WILD** | **A) ERRORS:** N = 442 obs, 380 ind  (56 ind – 2 trials, 10 ind – 3 trials) | | | | | **B) LATENCY:** N = 439 obs, 377 ind  (56 ind – 2 trials, 10 ind – 3 trials) | | | | |
| --- | --- | --- | --- | --- | --- | --- | --- | --- | --- | --- |
| *Fixed effects* | Median | Mean | *CI* | | pdirection | Median | Mean | *CI* | | pdirection |
| Intercept | **2.28** | **2.28** | **0.48** | **4.11** | **0.99** | **2.50** | **2.49** | **0.39** | **4.64** | **0.99** |
| ISA | -0.40 | -0.40 | -1.00 | 0.26 | 0.91 | 0.14 | 0.14 | -0.62 | 0.88 | 0.67 |
| Sex (male) | 0.09 | 0.09 | -0.14 | 0.32 | 0.78 | 0.09 | 0.09 | -0.16 | 0.34 | 0.76 |
| Age (juvenile) | 0.04 | 0.04 | -0.21 | 0.29 | 0.62 | 0.08 | 0.08 | -0.20 | 0.35 | 0.70 |
| Julian date | 0.00 | 0.00 | -0.01 | 0.01 | 0.63 | 0.00 | 0.00 | -0.01 | 0.01 | 0.55 |
| Time of day | -0.03 | -0.03 | -0.09 | 0.03 | 0.82 | -0.04 | -0.04 | -0.11 | 0.04 | 0.83 |
| Year (2022) | -0.07 | -0.07 | -0.32 | 0.18 | 0.70 | -0.09 | -0.09 | -0.37 | 0.19 | 0.73 |
| Year (2023) | -0.07 | -0.07 | -0.38 | 0.23 | 0.69 | -0.01 | -0.01 | -0.35 | 0.32 | 0.52 |
| Blood (yes) | -0.20 | -0.20 | -0.55 | 0.14 | 0.87 | -0.23 | -0.23 | -0.63 | 0.15 | 0.88 |
| Trial (2) | -0.16 | -0.16 | -0.59 | 0.28 | 0.76 | 0.10 | 0.10 | -0.29 | 0.50 | 0.69 |
| Trial (3) | -0.29 | -0.29 | -1.24 | 0.69 | 0.72 | -0.57 | -0.57 | -1.39 | 0.27 | 0.91 |
| ISA*Trial (2) | **1.24** | **1.24** | **0.34** | **2.14** | **1.00** |  |  |  |  |  |
| ISA*Trial (3) | -0.14 | -0.12 | -2.18 | 1.94 | 0.55 |  |  |  |  |  |
| *Random effects* |  |  |  |  |  |  |  |  |  |  |
| Individual ID (V_I_) | **0.49** | **0.49** | **0.31** | **0.69** |  | 0.28 | 0.29 | 0.00 | 0.61 |  |
| Site ID (V_SITE_) | 0.02 | 0.05 | 0.00 | 0.18 |  | 0.04 | 0.07 | 0.00 | 0.26 |  |
| Residual (V_R_) | **0.67** | **0.70** | **0.46** | **0.95** |  | **1.21** | **1.22** | **0.85** | **1.60** |  |
| Repeatability (*R*) | **0.4** | **0.4** | **0.25** | **0.55** |  | 0.18 | 0.19 | 0.00 | 0.38 |  |
| **2) COMMON GARDEN** | **A) ERRORS:** N = 153 obs, 72 ind  (54 ind – 2 trials, 32 ind – 3 trials) | | | | | **B) LATENCY:** N = 153 obs, 72 ind  (54 ind – 2 trials, 32 ind – 3 trials) | | | | |
| *Fixed effects* | Median | Mean | *CI* | | pdirection | Median | Mean | *CI* | | pdirection |
| Intercept | 1.05 | 1.05 | -0.78 | 2.96 | 0.86 | 3.04 | 3.01 | -0.46 | 6.73 | 0.95 |
| ISA | -0.21 | -0.22 | -0.99 | 0.57 | 0.71 | -0.02 | -0.02 | -1.14 | 1.14 | 0.52 |
| Sex (male) | 0.20 | 0.20 | -0.22 | 0.64 | 0.82 | 0.30 | 0.31 | -0.41 | 1.00 | 0.81 |
| Time of day | 0.06 | 0.06 | -0.12 | 0.24 | 0.74 | 0.00 | 0.00 | -0.35 | 0.35 | 0.50 |
| Trial (2) | -0.28 | -0.28 | -0.70 | 0.17 | 0.90 | 0.13 | 0.13 | -0.73 | 1.05 | 0.63 |
| Trial (3) | -0.43 | -0.43 | -0.87 | 0.02 | 0.97 | **-1.41** | **-1.43** | **-2.28** | **-0.54** | **1.00** |
| *Random effects* |  |  |  |  |  |  |  |  |  |  |
| Individual ID (V_I_) | 0.21 | 0.23 | 0.00 | 0.51 |  | 0.13 | 0.21 | 0.00 | 0.69 |  |
| Origin nest ID (V_NO_) | 0.04 | 0.08 | 0.00 | 0.28 |  | 0.04 | 0.09 | 0.00 | 0.55 |  |
| Foster nest ID (V_NF_) | 0.03 | 0.08 | 0.00 | 0.33 |  | 0.05 | 0.12 | 0.00 | 0.50 |  |
| Aviary ID (V_AV_) | 0.11 | 0.18 | 0.00 | 0.57 |  | 0.31 | 0.50 | 0.00 | 1.58 |  |
| Residual (V_R_) | **0.63** | **0.63** | **0.36** | **0.87** |  | **2.66** | **2.73** | **1.80** | **3.80** |  |
| Repeatability (*R*) | 0.35 | 0.37 | 0.06 | 0.71 |  | 0.22 | 0.24 | 0.02 | 0.51 |  |

**Table S3.** Model comparison to Table 1.1 and Table S2.1 when only including the first trial for wild birds. Posterior median and mean, credible intervals (CI), and pdirection (probability effect is in the same direction as median) for fixed and random effects in models using the 1) habitat effect and 2) ISA effect while evaluating the A) number of errors and B) latency to escape in a motor detour task. The number of errors was fit with negative binomial generalized linear mixed-effect models and the latency to escape was fit with a truncated lognormal mixed-effect models. The number of observations (obs), individuals (ind) are shown for each context and trait. Estimates whose credible intervals do not cross 0 are bolded.

| **1) Habitat** | **A) ERRORS:** N = 442 obs, 380 ind  (60 ind – 2 trials, 10 ind – 3 trials) | | | | | **B) LATENCY:** N = 439 obs, 377 ind  (56 ind – 2 trials, 10 ind – 3 trials) | | | | |
| --- | --- | --- | --- | --- | --- | --- | --- | --- | --- | --- |
| *Fixed effects* | Median | Mean | *CI* | | pdirection | Median | Mean | *CI* | | pdirection |
| Intercept | 1.82 | 1.79 | -0.41 | 3.91 | 0.94 | 1.89 | 1.90 | -0.38 | 4.39 | 0.94 |
| habitat | -0.25 | -0.24 | -0.91 | 0.45 | 0.82 | 0.09 | 0.10 | -0.76 | 0.90 | 0.61 |
| Sex (male) | 0.07 | 0.07 | -0.16 | 0.30 | 0.72 | 0.11 | 0.11 | -0.15 | 0.39 | 0.80 |
| Age (juvenile) | 0.01 | 0.01 | -0.24 | 0.26 | 0.54 | 0.06 | 0.06 | -0.21 | 0.37 | 0.66 |
| Julian date | 0.01 | 0.01 | 0.00 | 0.02 | 0.89 | 0.00 | 0.00 | -0.01 | 0.02 | 0.69 |
| Time of day | -0.02 | -0.02 | -0.08 | 0.05 | 0.71 | -0.02 | -0.02 | -0.10 | 0.06 | 0.68 |
| Year (2022) | -0.03 | -0.03 | -0.28 | 0.23 | 0.58 | -0.04 | -0.04 | -0.31 | 0.28 | 0.61 |
| Year (2023) | -0.16 | -0.16 | -0.46 | 0.14 | 0.86 | -0.01 | -0.01 | -0.37 | 0.36 | 0.53 |
| Blood (yes) | -0.38 | -0.38 | -0.79 | 0.01 | 0.97 | -0.24 | -0.24 | -0.70 | 0.24 | 0.83 |
| *Random effects* |  |  |  |  |  |  |  |  |  |  |
| Site ID (V_SITE_) | 0.05 | 0.09 | 0.00 | 0.28 |  | 0.05 | 0.08 | 0.00 | 0.29 |  |
| Residual (V_R_) | **1.59** | **1.60** | **1.26** | **1.96** |  | **1.5** | **1.5** | **1.27** | **1.75** |  |
| **2) ISA** |  |  |  | |  |  |  |  | |  |
| *Fixed effects* | Median | Mean | *CI* | | pdirection | Median | Mean | *CI* | | pdirection |
| Intercept | 1.92 | 1.92 | -0.15 | 3.94 | 0.97 | 1.72 | 1.73 | -0.75 | 4.07 | 0.92 |
| ISA | -0.43 | -0.42 | -1.02 | 0.24 | 0.92 | 0.14 | 0.14 | -0.53 | 0.86 | 0.70 |
| Sex (male) | 0.06 | 0.07 | -0.16 | 0.29 | 0.71 | 0.11 | 0.11 | -0.13 | 0.38 | 0.81 |
| Age (juvenile) | 0.02 | 0.02 | -0.22 | 0.25 | 0.56 | 0.07 | 0.07 | -0.22 | 0.35 | 0.68 |
| Julian date | 0.01 | 0.01 | -0.01 | 0.02 | 0.85 | 0.00 | 0.00 | -0.01 | 0.02 | 0.73 |
| Time of day | -0.02 | -0.02 | -0.08 | 0.05 | 0.69 | -0.02 | -0.02 | -0.10 | 0.06 | 0.66 |
| Year (2022) | -0.02 | -0.02 | -0.27 | 0.24 | 0.55 | -0.04 | -0.04 | -0.32 | 0.28 | 0.60 |
| Year (2023) | -0.15 | -0.15 | -0.44 | 0.15 | 0.84 | -0.01 | -0.01 | -0.35 | 0.36 | 0.53 |
| Blood (yes) | -0.37 | -0.38 | -0.79 | 0.01 | 0.97 | -0.23 | -0.23 | -0.69 | 0.25 | 0.83 |
| *Random effects* |  |  |  |  |  |  |  |  |  |  |
| Site ID (V_SITE_) | 0.02 | 0.05 | 0.00 | 0.19 |  | 0.04 | 0.08 | 0.00 | 0.29 |  |
| Residual (V_R_) | **1.58** | **1.59** | **1.26** | **1.94** |  | **1.50** | **1.50** | **1.26** | **1.75** |  |

**Table S4.** Model comparison to Table 1.1 in main text when using a quantitative genetic animal model to estimate heritability in the wild. Posterior median and mean, credible intervals (CI), and pdirection (probability effect is in the same direction as median) for fixed and random effects evaluating the A) number of errors and B) latency to escape in a motor detour task. The number of errors was fit with negative binomial generalized linear mixed-effect models and the latency to escape was fit with truncated lognormal mixed-effect models. The number of observations (obs), individuals (ind), and individual repeated individual measures are shown for each context and trait. Estimates whose credible intervals do not cross 0 are bolded.

| **WILD** | **A) ERRORS:** N = 424 obs. 362 ind  (56 ind – 2 trials. 10 ind – 3 trials) | | | | | **B) LATENCY:** N = 422 obs. 360 ind  (56 ind – 2 trials. 10 ind – 3 trials) | | | | |
| --- | --- | --- | --- | --- | --- | --- | --- | --- | --- | --- |
| *Fixed effects* | Median | Mean | *CI* | | pdirection | Median | Mean | *CI* | | pdirection |
| Intercept | 1.65 | 1.64 | -0.26 | 3.54 | 0.96 | 2.14 | 2.12 | -0.09 | 4.33 | 0.97 |
| ISA | -0.16 | -0.16 | -0.73 | 0.40 | 0.77 | 0.17 | 0.17 | -0.49 | 0.83 | 0.75 |
| Sex (male) | 0.06 | 0.06 | -0.17 | 0.30 | 0.69 | 0.05 | 0.05 | -0.21 | 0.30 | 0.65 |
| Age (juvenile) | 0.06 | 0.06 | -0.20 | 0.31 | 0.66 | 0.07 | 0.07 | -0.21 | 0.35 | 0.68 |
| Julian date | 0.01 | 0.01 | 0.00 | 0.02 | 0.89 | 0.00 | 0.00 | -0.01 | 0.02 | 0.64 |
| Time of day | -0.03 | -0.02 | -0.09 | 0.04 | 0.77 | -0.03 | -0.03 | -0.10 | 0.05 | 0.77 |
| Year (2022) | -0.15 | -0.15 | -0.41 | 0.10 | 0.88 | -0.12 | -0.12 | -0.41 | 0.16 | 0.80 |
| Year (2023) | -0.11 | -0.11 | -0.42 | 0.20 | 0.76 | -0.02 | -0.02 | -0.36 | 0.33 | 0.54 |
| Blood (yes) | -0.30 | -0.30 | -0.63 | 0.04 | 0.96 | -0.25 | -0.25 | -0.62 | 0.13 | 0.90 |
| Trial (2) | 0.24 | 0.24 | -0.11 | 0.59 | 0.91 | 0.11 | 0.11 | -0.28 | 0.51 | 0.72 |
| Trial (3) | -0.33 | -0.32 | -1.06 | 0.43 | 0.80 | -0.59 | -0.59 | -1.40 | 0.24 | 0.92 |
| ISA*Trial (2) | 1.65 | 1.64 | -0.26 | 3.54 | 0.96 | 2.14 | 2.12 | -0.09 | 4.33 | 0.97 |
| ISA*Trial (3) | -0.16 | -0.16 | -0.73 | 0.40 | 0.77 | 0.17 | 0.17 | -0.49 | 0.83 | 0.75 |
| *Random effects* |  |  |  |  |  |  |  |  |  |  |
| Pedigree (V_A_) | 0.35 | 0.34 | 0.00 | 0.57 |  | 0.26 | 0.27 | 0.00 | 0.60 |  |
| Individual ID (V_PE_) | 0.09 | 0.14 | 0.00 | 0.43 |  | 0.08 | 0.14 | 0.00 | 0.44 |  |
| Site ID (V_SITE_) | 0.02 | 0.05 | 0.00 | 0.27 |  | 0.04 | 0.07 | 0.00 | 0.26 |  |
| Residual (V_R_) | **0.73** | **0.74** | **0.51** | **0.99** |  |  |  |  |  |  |
| heritability (*h^2^*) | 0.28 | 0.27 | 0.00 | 0.45 |  | 0.16 | 0.17 | 0.00 | 0.37 |  |

**Table S5.** Posterior median and mean, credible intervals (CI) for fixed and random effects evaluating the association between the five significant SNPs and number of errors in a motor detour task for wild birds. The number of errors was fit with negative binomial generalized linear mixed-effect models. Fixed- and random- effect estimates whose credible intervals do not cross 0 or are greater than 0.001, respectively, are bolded. The five SNPs fitted as fixed effects were the ones that were significant in the GWAS analysis. Note that the sum of the individual SNP effects does not equal the total SNP effect because the sum of the medians differs from the median of the sums.

| *Fixed effects* | Median | Mean | | CI | |
| --- | --- | --- | --- | --- | --- |
| Intercept | **0.97** | **0.97** | | **0.44** | **1.48** |
| SNP1 (Chrom 8: gene unknown) | **0.26** | **0.26** | | **0.07** | **0.45** |
| SNP2 (Chrom 13 gene: LOC107210618/ LOC117245099) | **0.55** | **0.55** | | **0.32** | **0.77** |
| SNP3 (Chrom 14 gene: SYNGR3) | **-0.53** | **-0.53** | | **-0.82** | **-0.24** |
| SNP4 (Chrom 23: gene unknown) | **-0.22** | **-0.22** | | **-0.40** | **-0.05** |
| SNP5 (Chrom 24 gene: POU2AF2/ HINFP) | **0.31** | **0.31** | | **0.14** | **0.48** |
| *Random effects* |  |  |  | |  |
| SNP1 | 0.03 | 0.03 | 0.00 | | 0.08 |
| SNP2 | **0.10** | **0.10** | **0.03** | | **0.19** |
| SNP3 | **0.06** | **0.06** | **0.01** | | **0.13** |
| SNP4 | 0.03 | 0.03 | 0.00 | | 0.07 |
| SNP5 | 0.06 | 0.06 | 0.00 | | 0.12 |
| Hmatrix (V_A__nonSNP) | 0.10 | 0.13 | 0.00 | | 0.34 |
| Individual ID (V_PE_) | 0.18 | 0.18 | 0.00 | | 0.39 |
| Site ID (V_SITE_) | 0.07 | 0.11 | 0.00 | | 0.36 |
| Residual (V_R_) | **0.70** | **0.71** | **0.48** | | **0.97** |
| *Proportion variation explained* |  |  |  | |  |
| Total SNPs | **0.21** | **0.21** | **0.12** | | **0.30** |
| SNP1 | 0.02 | 0.02 | 0.00 | | 0.05 |
| SNP2 | **0.07** | **0.07** | | **0.02** | **0.13** |
| SNP3 | 0.04 | 0.05 | | 0.00 | 0.09 |
| SNP4 | 0.02 | 0.02 | | 0.00 | 0.05 |
| SNP5 | 0.04 | 0.04 | | 0.00 | 0.09 |
| Individual ID (V_PE_) | 0.13 | 0.13 | | 0.00 | 0.28 |
| Site ID (V_SITE_) | 0.05 | 0.07 | | 0.00 | 0.22 |
| Residual (V_R_) | **0.50** | **0.50** | | **0.35** | **0.65** |
| Heritability (*h^2^_nonSNP*) | 0.07 | 0.09 | | 0.00 | 0.24 |
